# Control of cortical population activity with patterned microstimulation

**DOI:** 10.64898/2026.03.02.709018

**Authors:** Giacomo Barzon, Anandita De, Isaac Moran, Conner Carnahan, Luca Mazzucato, Roozbeh Kiani

**Author notes:** co-first author (Giacomo Barzon, Anandita De, Isaac Moran). co-senior author.

## Abstract

Closed-loop control of cortical activity is a central goal in systems neuroscience and clinical neuromodulation, but most approaches either rely on detailed circuit models that are unattainable in vivo or on open-loop stimulation tuned by trial and error. Here we introduce REACHable manifold Control (REACH-Ctrl), a data-driven brain–computer interface that achieves real-time control of population spiking activity using patterned microstimulation and multi-electrode recordings. REACH-Ctrl learns a finite-horizon controllability map directly from short training epochs in which random multi-electrode pulse sequences are delivered through a subset of electrodes while recording evoked responses. From these input-output data, it identifies the “reachable manifold” of population states and computes low-current microstimulation sequences that steer activity toward designated targets, without explicit knowledge of the underlying connectivity or dynamics. We test REACH-Ctrl in macaque prefrontal cortex, demonstrating high control accuracy, robust across sessions and stimulation parameters. Geometric analyses showed that multi-pulse sequences traverse a well-defined reachable manifold with substantial, but incomplete, overlap with the intrinsic neural activity manifold, revealing both on- and off-manifold components of control. Encoding models further revealed that, in our weak-stimulation regime, population responses to multi-electrode sequences are well approximated by the linear sum of localized “stimulation fields” with modest history dependence, explaining the success of our linear control approach. These results demonstrate precise, sample-efficient control of cortical population activity with clinically relevant microstimulation hardware, and provide a general blueprint for designing perturbations for sparsely observed neural circuits.

## Introduction

The ability to directly perturb the brain to achieve a desired target effect using electrical stimulation holds broad therapeutic promise across neurological and psychiatric disorders [1–9]. Perturbation experiments have led to important insights into the organization and function of neural circuits in the primate [10–12] and rodent brain [13–16]. Yet most studies rely on open-loop interventions, and stimulation parameters are typically chosen without predictive models that link stimulation patterns to expected population responses, forcing reliance on inefficient trial-and-error (with exceptions [17]).

Control-theoretic frameworks offer a route to principled perturbation design in complex networks [18]. Recent work has used structural and functional imaging to study controllability of large-scale brain dynamics in humans [19–22]) and in inter-areal stimulation experiments in non-human primates [23]. However, it is not known how to achieve spatiotemporally precise control of neural-population spiking activity within a circuit. Current approaches typically assume access to a dynamical model or structural connectome for analyzing how inputs shape network trajectories. But this approach faces both conceptual and practical barriers. Conceptually, inferring a high-dimensional recurrent network from sparse, noisy recordings is a challenging inverse problem: many distinct dynamical systems can reproduce the same observed trajectories under limited sampling [24, 25]. Practically, full connectomes cannot be measured in behaving subjects, and current recording technologies sample only a tiny fraction of neurons in any given area. These constraints limit model-based control strategies that rely on explicit system identification of connectivity or dynamics [19–22, 26].

At the same time, experimental work on causal control of local circuits has largely focused on optical and optogenetic methods with single-cell or even subcellular precision [27, 28]. Two-photon imaging and patterned photostimulation provide powerful tools to probe and manipulate neurons in rodents [13–15, 28, 29], and have transformed our understanding of circuit mechanisms. However, these approaches require optical access and genetic tools that are not currently available for routine use in humans and face substantial practical and ethical barriers for widespread clinical deployment [30–34]. In contrast, electrical microstimulation and related approaches (including deep brain stimulation) lack single-neuron and cell-type specificity [35, 36], but are already used in humans and are widely deployed in both clinical and research settings [1–3, 37]. A central open question is whether such coarse, clinically feasible stimulation can nonetheless be harnessed to achieve precise control of high-dimensional population activity, and if so, how to design stimulation protocols that are both data-efficient and robust in the face of unknown circuit dynamics.

Here, we bridge this gap by implementing a closed-loop brain–computer interface (BCI) in macaque prefrontal cortex that achieves real-time control of neural population spiking activity using patterned microstimulation and multi-electrode recordings (Fig. 1a). We introduce REACHable manifold Control (REACH-Ctrl), an algorithm that operates directly on input-output data. In each session, REACH-Ctrl delivers short sequences of random multielectrode pulses through a subset of electrodes while recording the evoked responses. From these stimulation-response data, it identifies the set of population activity patterns that can be evoked by microstimulation, and designs optimal microstimulation sequences that steer activity toward target patterns with minimal current injection (Fig. 1b-d). REACH-Ctrl builds on recent advances in data-driven control [38–41], bypassing explicit system identification.

**FIG. 1.**
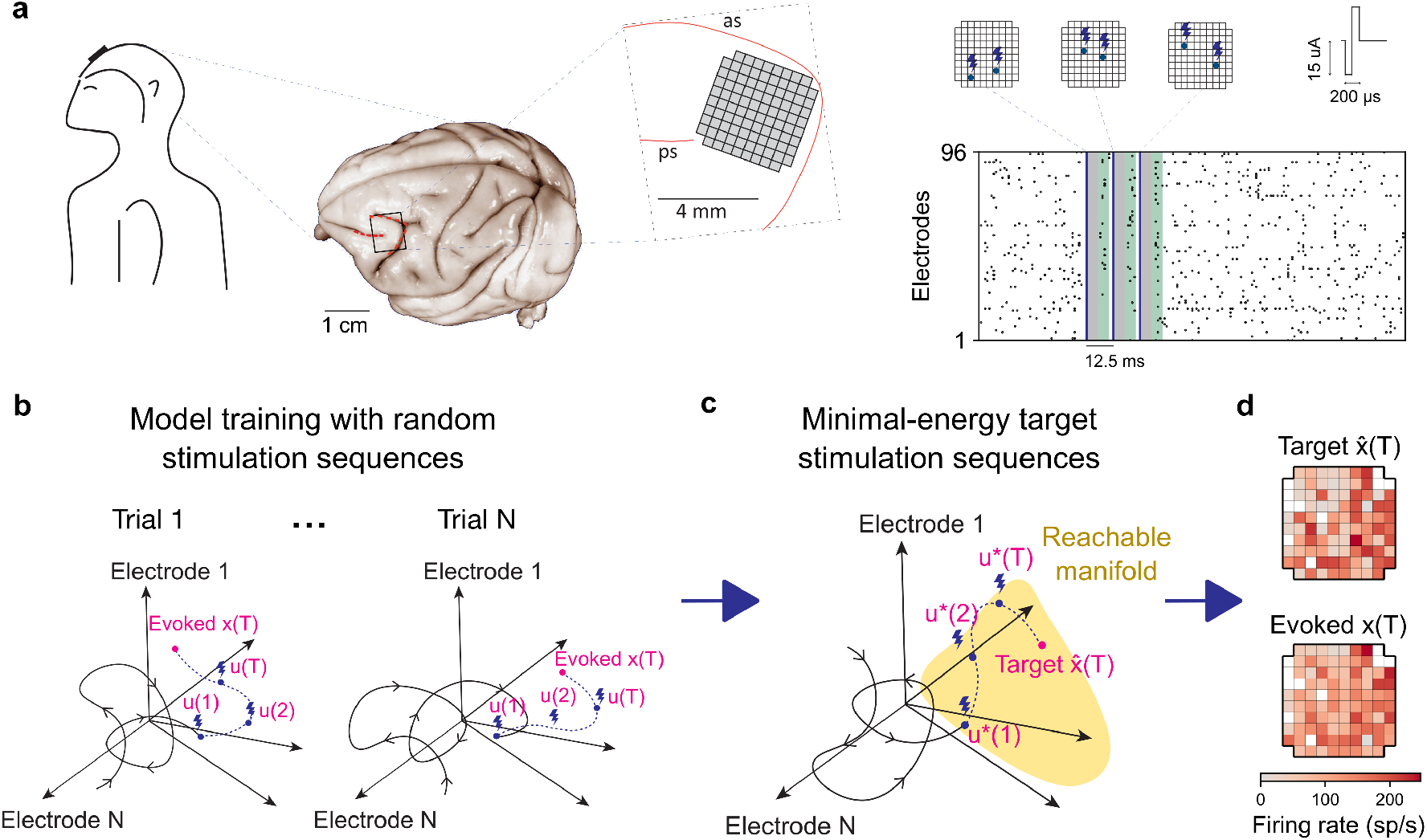
A brain-computer interface for control of cortical population activity. **a)** Experimental setup and neural recording. Left: a 96-channel Utah array chronically implanted in the pre-arcuate gyrus of macaque monkeys. Center: anatomical location of the array relative to the arcuate sulcus (as) and principal sulcus (ps). Brain picture adopted from the University of Wisconsin Brain Collection. Right: example multi-electrode microstimulation sequences and the resulting population spiking activity recorded across the electrodes. Evoked responses were quantified in a 5 ms window (green); earlier times were excluded (gray) to allow hardware switching between stimulation mode and artifact-free recording. **b)** In the training epoch, REACH-Ctrl learns stimulus-response mapping by delivering random T-pulse stimulation sequences u(1:T) while recording the evoked population responses x(1:T). **c)** Using training data, REACH-Ctrl estimates the “reachable manifold” (yellow), representing the set of neural activity patterns reachable via microstimulation. Given a reachable target pattern 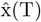 (T), REACH-Ctrl computes the stimulation sequence u^*^(1:T) required to steer the ensemble activity toward the target. **d)** Comparison between a target activity pattern 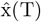 (T) and the population pattern x(T) evoked by u^*^(1:T). Each cell in 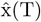 (T) and x(T) represents one electrode.

We tested REACH-Ctrl using chronically implanted Utah arrays in behaving monkeys (Fig. 1a), a configuration that closely parallels invasive BCIs already used in humans [37, 42]. Within single sessions, the algorithm learns the reachable manifold from short training epochs and then generates microstimulation sequences that evoke target population patterns with high accuracy. REACH-Ctrl is robust to noise and sparse sampling, and it achieves target population patterns precisely. Geometric analyses show that multi-pulse sequences traverse a well-defined reachable manifold with substantial, but incomplete, overlap with the intrinsic neural manifold defined by pre-stimulation activity. This reveals both on- and off-manifold components of control and provides a principled way to visualize how stimulation-driven trajectories explore the space of reachable population states. Finally, we use encoding models to characterize “stimulation fields”—the spatiotemporal mapping from multi-electrode pulse sequences to population responses—and to uncover the compositional rules underlying multi-electrode stimulation sequences. In the weak-stimulation regime we adopt, responses to multi-electrode sequences are well approximated by the linear sum of localized stimulation fields with modest history dependence, and nonlinear interaction terms contribute only small additional variance. Together with the manifold geometry, these results explain the success of our data-driven linear control approach and delineate its operating range. More broadly, our findings demonstrate precise, sample-efficient control of cortical population activity using clinically relevant microstimulation hardware in single experimental sessions, and establish a general blueprint for designing perturbations in sparsely observed neural circuits *in vivo*.

## Results

We chronically implanted 96-channel Utah arrays in the pre-arcuate gyrus (Fig. 1a) and used the same electrodes for both recording and microstimulation during quiet wakefulness (n = 30 sessions in 2 monkeys). Electrodes were spaced at 400 µm, providing approximately one electrode per cortical microcolumn and sampling a 4×4 mm^2^ patch of cortex. To control population activity in this patch, we developed REACH-Ctrl, a data-driven approach suitable for fast and efficient control of evoked population activity patterns in single sessions (Fig. 1b-d).

REACH-Ctrl steers ensemble spiking activity x(t) on the array towards a target pattern 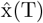 (T) over T stimulation steps (pulses). For a desired target state 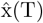 (T), REACH-Ctrl computes an optimal sequence of microstimulation pulses u^*^(1:T), where each u(t) is an m-dimensional binary vector indicating the subset of electrodes stimulated at step t.Each session comprised a training block followed by a test block (Fig. 1b-d). During the training block, we delivered randomized multi-electrode stimulation sequences and recorded the resulting population responses (Fig. 1b). From these input–output data we identified the subspace of reachable targets (i.e., the reachable manifold) and computed optimal stimulation sequences to drive the population toward selected targets with a small number of pulses (Fig. 1c,d).

### A data-driven controller for cortical population activity

The first phase of REACH-Ctrl involved collecting stimulation–response pairs to train the model. We selected a subset of m electrodes for stimulation based on their high functional connectivity [43]. Each stimulation sequence comprised T pulses with inter-pulse interval Δt, each pulse simultaneously stimulating a random subset κ of the chosen m electrodes (m = 10, T = 3, κ = 2, Δt = 12.5 ms in 30 standard sessions; parameters were systematically varied in 12 additional sessions). The training epoch in standard sessions consisted of N_train_ = 100 T-pulse microstimulation trials. The sample size was chosen to satisfy the “rank condition” N_train_ > mT, required for the estimation of the set of reachable patterns (see Methods). We restricted our protocol to weak microstimulation currents (15 µA), below the thresholds that evoked eye movements or directly activated cells outside the target microcolumns [35, 36, 44–46].

Between pulses, the BCI toggled from stimulation to recording mode, enabling us to record spiking responses immediately following each pulse (Fig. 2a). We restricted analyses to electrodes with low noise and clear action potential waveforms (n = 90 for Monkey 1, n = 86 for Monkey 2). Training data were compiled into a binary mT×N_train_ input matrix U, encoding the stimulated electrodes across pulses in each trial, and an nT× N_train_ response matrix X, encoding population responses to each pulse (5 ms bins); the bottom n rows of X encoded the response at the end of the microstimulation sequence. The training data U, X supported the online determination of reachable targets and design of the optimal controller, and offline analyses of stimulation fields (see Methods).

**FIG. 2.**
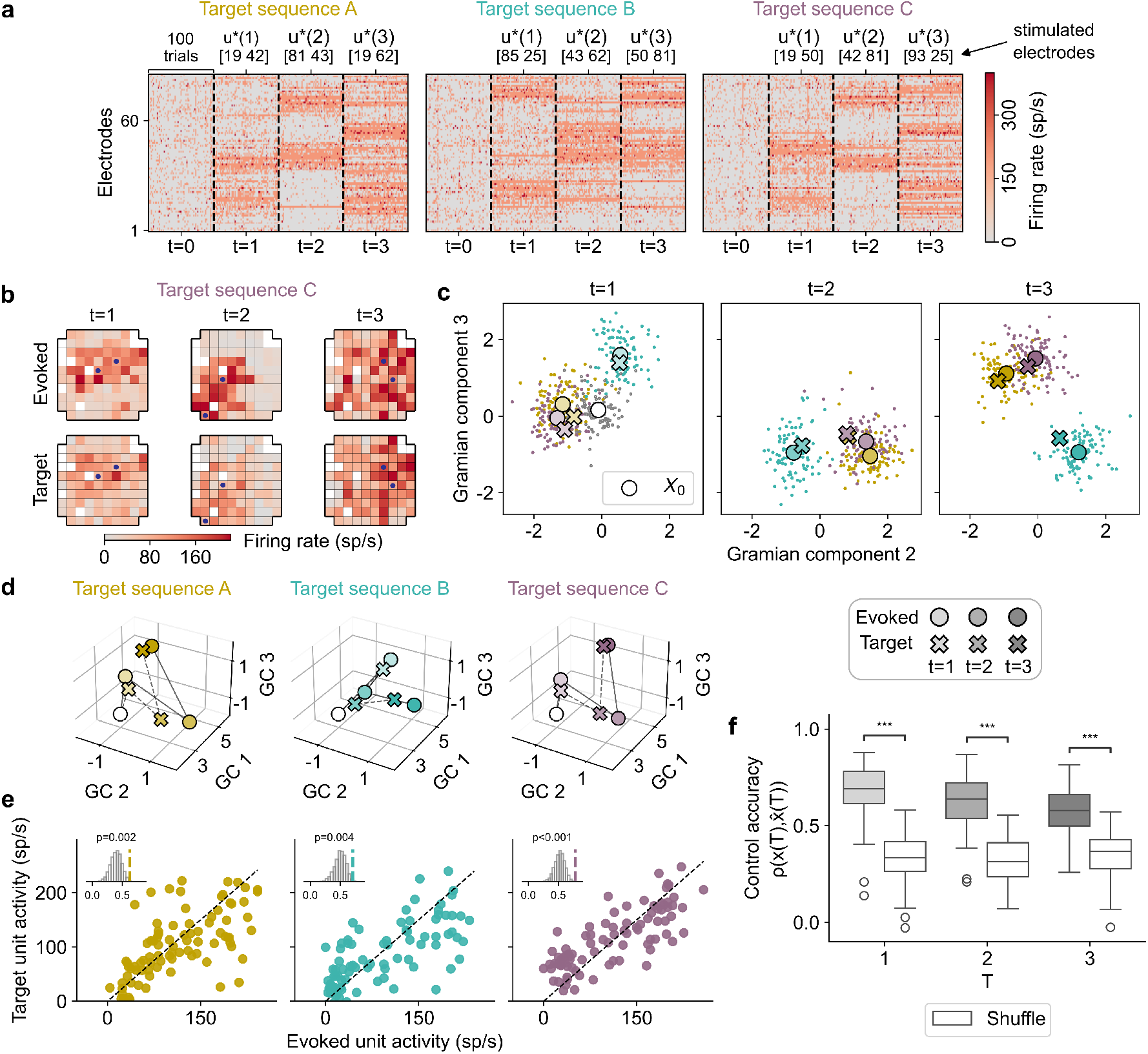
REACH-Ctrl steers population spiking activity toward designated targets. **a)** Representative T = 3 stimulation sequences and ensemble responses during a test session (yellow, teal, and purple denote sequences A, B, and C, respectively). Each panel shows the pre-stimulation state (t = 0) and responses to N_test_ = 100 repetitions of each pulse u^*^(t = 1, 2, 3); trials are juxtaposed horizontally at each time step). **b)** Comparison of evoked and target population patterns for example sequence C. Spatial heatmaps show the trial-averaged evoked activity (top) and the predicted target activity (bottom) at each time step. **c)** Evoked (circles) and target population activity (crosses) projected onto the second and third Gramian Components (GC; see Methods) exhibit tight alignment at each time step for sequences A, B, C. Dots: single trials. Colors as in panel a. **d)** Evoked and target neural trajectories in three-dimensional GC space for sequences A, B, C. **e)** Trial-averaged evoked firing rates versus target firing rates for all units at the final time step (T = 3) for the three sequences reveal high control accuracy, defined as the Pearson correlation *ρ*(x(T), 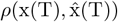 (T)). Insets: empirical *ρ* compared to shuffled distributions. **f)** Distribution of control accuracy across sequences of all sessions at each time step. Grey and while boxplots show empirical and shuffled values. ***: p *<* 0.001.

At the end of the training epoch, we used REACH-Ctrl to determine the set of population activity patterns that could be reached by microstimulation (the reachable manifold in Fig. 1c; see Methods). Our goal was to steer the population response x(t) to a designated target 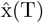 (T) in T steps. REACH-Ctrl is rooted in Willems’ Fundamental Lemma [47], which demonstrates that any reachable population trajectory can be expressed as linear combinations of trajectories observed during the training epoch. REACH-Ctrl determines the set of reachable patterns as the span of the columns of the controllability matrix 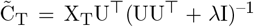. The ridge regularization λ was included to improve robustness and was optimized online via cross-validation (Supplementary Fig. S2). We selected target patterns 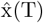 (T) within this reachable manifold and, for each of them, used REACH-Ctrl to estimate the corresponding optimal sequence u^*^(1 : T) to evoke them (Fig. 2).

Reachable manifold, target patterns, and optimal stimulation sequences were computed online at the end of the training block. During the subsequent test block, each optimal sequence was delivered for N_test_ = 100 interleaved trials (3-4 distinct targets per session; Fig. 2a). As in training, the BCI switched between stimulation and recording after each pulse, allowing us to compare evoked responses with target patterns (Fig. 2b-f).

### REACH-Ctrl steers population activity toward designated target patterns

We first examined whether the terminal responses x(T) evoked by each optimal sequence u^*^(1 : T) matched the predicted targets 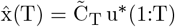. To visualize REACH-Ctrl performance, we projected population activity onto the top Gramian Component Analysis axes, which captured the most easily reachable directions. Evoked activity patterns closely matched the target patterns, and this alignment occurred not only in response to the last pulse x(T), but at all steps in the sequence x(1:T) (Fig. 2b-d). The target patterns were elaborate; instead of a simple gain on population activity, they involved nuanced patterns with distinct firing rates across the array. We found a high and significant control accuracy across sessions, quantified as the Pearson correlation between evoked and target final states, 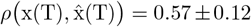 (mean±s.d.) across test sequences (Fig. 2f; significantly above shuffled surrogates, 0.35 ± 0.11, p < 10^−5^, obtained by permuting the columns of the data matrix X_T_). Comparing the time course of target patterns 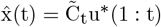 and evoked responses x(t) after each pulse in the sequence confirmed a high control accuracy at each step (Fig. 2f), consistent with REACH-Ctrl accurately controlling the full temporal evolution of trajectories. A small but significant decrease of control accuracy was present along the time course (mixed linear model ρ = ρ_0_ + *β*t + b: ρ_0_ = 0.735± 0.017, mean± s.e.; *β* = –0.054± 0.007; random intercept var(b) = 0.008± 0.020, p < 0.001; see Methods), consistent with the theoretical expectation from data-driven control of noisy systems [48].Control accuracy was robust across experimental parameters. Comparable accuracy was obtained for pulse sequences optimized for different inter-pulse intervals (Δt = 12.5, 25, 50 ms) and for stimulating electrodes separated by different spatial distances on the array (Supplementary Fig. S2c,e). When we varied the number of simultaneously stimulated electrodes per pulse, control accuracy remained high and significant across the tested range of 1-3 electrodes and peaked for two electrodes (Supplementary Fig. S2d), supporting our choice of two electrodes per pulse in the standard sessions.

Microstimulation-evoked patterns were stimulation-specific (Fig. 2a-d). Activity patterns evoked by pulses within and across stimulation sequences could be reliably decoded with high accuracy at each pulse in the sequence (cross-validated Linear Discriminant Analysis, 0.93± 0.06, mean ± s.d.; Supplementary Fig. S3b-c). This discriminability exhibited a small but significant linear increase from the first to the third pulse in a sequence (mixed linear model, *β* = 0.028± 0.007, mean ± s.e., p < 0.001, Supplementary Fig. S3c), confirming a cumulative effect of sequential stimulations on the separation of neural states. Furthermore, while control performance remained robust, we found that discriminability between pairs of responses slightly decreased when the set of stimulated pulses featured overlapping sets of electrode (Supplementary Fig. S3d). This effect is consistent with a summation rule for the effects of multi-electrode stimulation, which we examine below.

### Sufficiency of short stimulation sequences

To probe how individual pulses contribute to control, we compared optimal three-step sequences u^*^ = [u_1_, u_2_, u_3_] to reduced two-step variants omitting either the first or second pulse (Fig. 3a). Reduced sequences still drove activity toward the target pattern 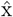 above chance (Fig. 3b), indicating that a single omission does not abolish the effects of the preserved pulses. However, omitting the second pulse caused a significantly larger deviation from the target than omitting the first (Fig. 3b). We next compared the representational geometry of responses evoked by reduced versus unaltered sequences. Altered sequences evoked activity patterns distinct from the unaltered sequence and from each other (average LDA classification accuracy, 0.65 ±0.06, mean± s.d.; Fig. 3c,d), demonstrating sensitivity to the exact stimulation sequence. Moreover, omission of the second pulse led to stronger discriminability from the unaltered sequence than omission of the first (LDA false-positive rate, 0.14± 0.07 vs. 0.25± 0.09; Fig. 3c,d).

**FIG. 3.**
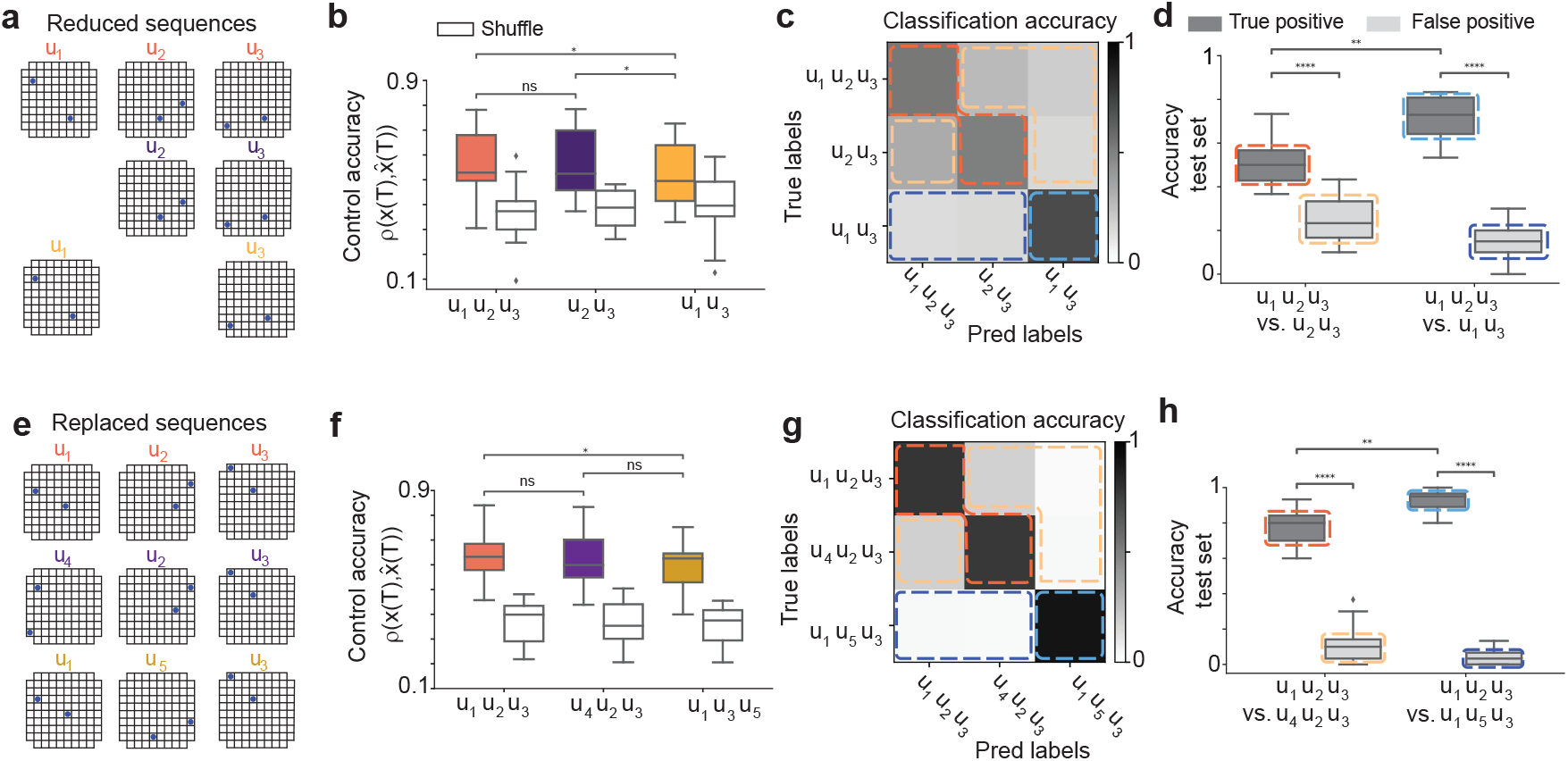
Sufficiency of short sequences. **a-d)** Altered stimulation sequences with one-pulse omission. (a) Schematic of stimulated electrodes on the array for an optimal sequence and altered variants in an example session. Top: full sequence; middle and bottom: sequences with the first or second pulse omitted. (b) Control accuracy, quantified as the Pearson correlation between the target 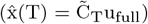 and evoked activity x(T) for full vs. reduced sequences. (c) Confusion matrix of evoked responses to full vs. altered sequences, quantified by a linear discriminant classifier trained to decode the evoked activity. (d) Comparison of true- and false-positive classification accuracy for full vs. reduced sequences across sessions. **e-h)** Altered sequences with pulse replacement. (e) Schematic of an optimal sequence (top) and altered sequences in which the first or second pulse is replaced by random pairs of stimulating electrodes (middle and bottom). (f-h) Same analyses as in panels a-d, but for pulse replacement rather than omission.

We further dissected pulse-specific contributions by replacing individual pulses in the optimal three-step sequences with random pulses (Fig. 3e). Both altered sequences still yielded significant control accuracy, and replacing the second pulse produced a larger drop in accuracy compared to the first pulse (Fig. 3f,h). Responses evoked by altered vs. unaltered sequences were discriminable, with second-pulse replacement producing stronger discriminability from the unaltered sequence than first-pulse replacement (Fig. 3g).

Together, these experiments indicate that multi-pulse microstimulation engages history-dependent effects, with later pulses exerting the strongest influence on the final activity. They also show that short sequences are sufficient for above-chance control of target patterns within the reachable subspace.

### Traversability of the reachable subspace

To assess how multi-step microstimulation reshapes population trajectories, we compared the geometry of stimulation-evoked activity to two manifolds in state space: the reachable manifold ℳ_reach_, defined by the column space of 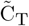, and the intrinsic manifold ℳ_0_, defined by the principal components of the n *×* N_test_-dimensional pre-stimulation activity X_0_ (Fig. 4a-c). This analysis allowed us to ask whether multi-pulse sequences simply pushed the circuit along intrinsic modes of variability or also drove it into directions that were rarely explored during ongoing activity. We used two complementary metrics: (i) the Euclidean distance from baseline activity (Fig. 4) and (ii) the fraction of variance of the reachable and intrinsic manifolds explained by the evoked trajectories (Fig. S4).

**FIG. 4.**
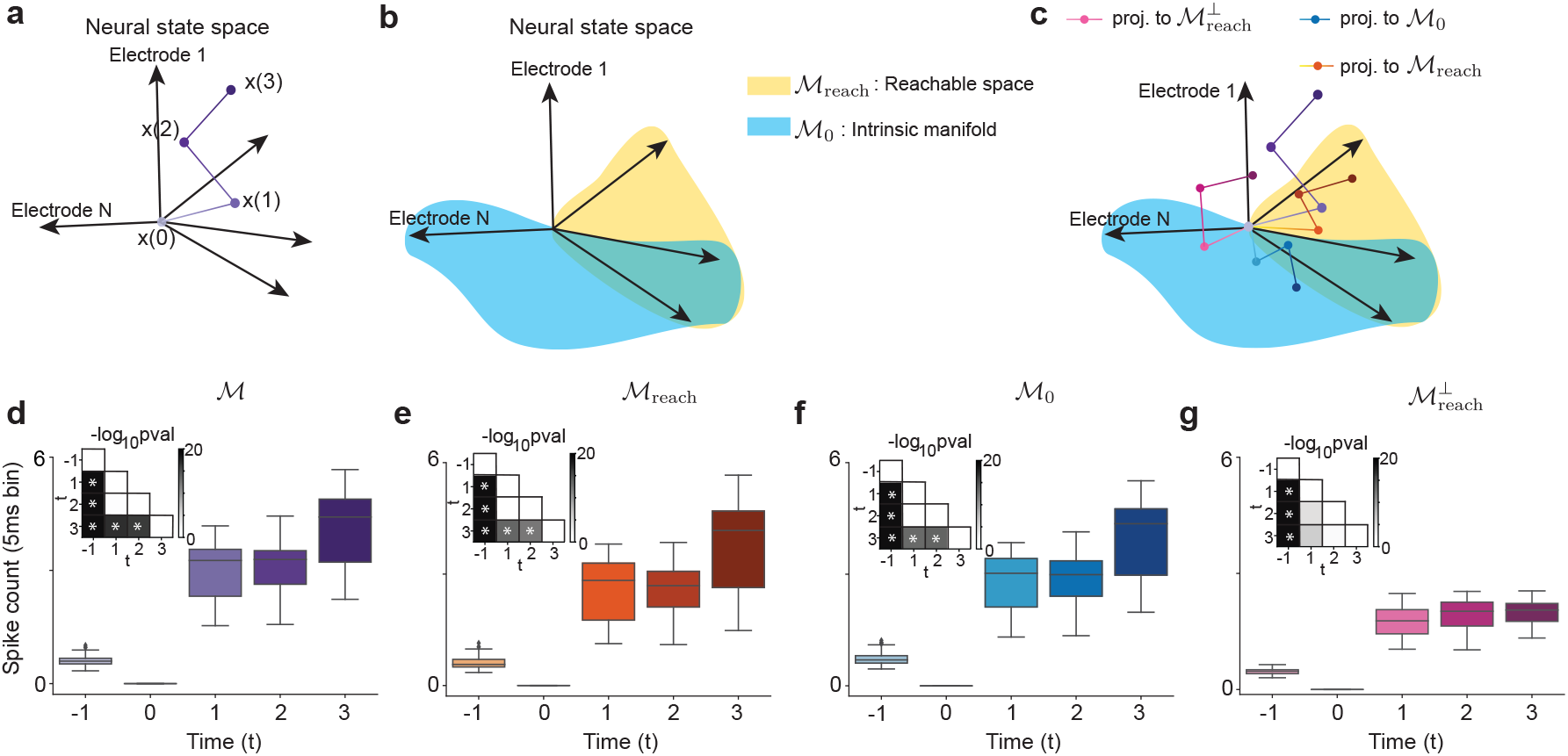
Traversability of reachable manifold. **a)** Schematic of the evoked trajectory x(t) in the full-dimensional data space. **b)** Schematic of the reachable manifold ℳ _reach_, the intrinsic manifold ℳ _0_, and their overlap ℳ _reach_ ∩ ℳ _0_. **c)** Schematic of the evoked trajectory projected onto different subspaces: full space (purple), reachable manifold ℳ _reach_ (orange), intrinsic manifold ℳ_0_ (blue), and the subspace orthogonal to the reachable manifold in the full space 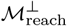 (magenta). **d-g)** Distance of the pre-stimulation state x(0) from evoked activity x(t) following each pulse (t *>* 0), as well as from spontaneous activity before stimulation start (t = –1). Boxplots show distribution of distances across sequences and sessions (5-ms bins). Insets show –log_10_p values from paired Wilcoxon tests comparing distance distributions at different times. (d) Distance of x(0) and x(t) in the full space. (e) Distance of projections onto the reachable manifold. (f) Distance of projections onto the intrinsic manifold. (g) Distance of projections onto the subspace orthogonal to the reachable manifold.

We first asked whether stimulation sequences drive trajectories away from baseline. The distance between the evoked state x(t) and the pre-stimulation state x(0) was significantly above baseline and increased with consecutive pulses (Fig. 4d). This distance exceeded the typical distance between baseline spike-count vectors (Fig. 4d), indicating that stimulation produces a substantial displacement of population spiking activity.

To identify where this displacement lies, we projected the evoked trajectories either onto the reachable manifold ℳ _reach_ or onto the intrinsic-activity manifold ℳ_0_. We then re-computed the distance-from-baseline metric in each projection. The progressive excursion with pulse number was preserved when trajectories were projected onto either ℳ _reach_ or ℳ _0_ (Fig. 4c-f). In contrast, this progressive excursion was absent in the component orthogonal to ℳ _reach_ (denoted 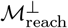), indicating that the cumulative effect of successive pulses is confined to the reachable subspace.

To dissect the relative role of ℳ _reach_ and ℳ _0_, we quantified their overlap. The fraction of variance of ℳ_reach_ accounted for by directions within ℳ_0_ was high (0.8 ± 0.02, mean±s.d.; see Methods and Fig. 4c,d), suggesting that the stimulation sequence kept activity largely within the intrinsic manifold. We then split the reachable manifold into a component overlapping with the intrinsic one, ℳ_reach_ *∩ℳ*_0_, and a complementary component, ℳ _reach_ \ℳ_0_. The progressive excursion away from baseline was present in both components (Supplementary Fig. S4b,c).

Together, these analyses show that progressing along the multi-step sequence pushes activity farther from baseline while keeping the displacement within ℳ_reach_. Within that subspace, trajectories follow directions that are shared with the intrinsic manifold as well as directions that are part of the reachable manifold but lie off the intrinsic manifold. We confirmed these conclusions with a complementary metric: the fraction of variance of each manifold explained by the evoked trajectories (Supplementary Fig. S4d-h). Trajectories explained a large fraction of variance within both ℳ _reach_ and ℳ _0_, as well as their shared subspace ℳ _reach_ ∩ ℳ _0_ (Supplementary Fig. S4d,e). Moreover, the variance explained within ℳ _reach_ increased with pulse number, consistent with progressively deeper engagement of the reachable manifold by multi-pulse sequences.

These results highlight two features of population responses to optimal multi-pulse sequences: (i) successive pulses progressively drive activity deeper into the reachable manifold of REACH-Ctrl, and (ii) this evolution concurrently follows intrinsic-manifold directions that are shared with the reachable manifold.

### Microstimulation effects sum linearly

To uncover the features that best explained neural responses to microstimulation, we fit cross-validated encoding models to training-epoch data and used them to predict test-epoch responses (Fig. 5a). Although Poisson Generalized Linear Models are standard for spike encoding [49, 50], our theoretical analysis showed that, in the weak microstimulation regime we adopted, these models reduce to equivalent linear forms (Methods and Supplementary Fig. S5). We therefore focused on linear models relating spiking activity to stimulation and spike history. We first compared a full model—which included stimulation history, spike history, and nonlinear interactions among stimulating electrodes and between stimulation and spike history—with single-feature models. Stimulation history alone captured nearly all the predictive power and outperformed models based on other features (Fig. 5b). Because collinearity can inflate the apparent explanatory power of single-feature models, we next assessed unique contributions by shuffling each feature while leaving the others intact [51, 52]. This analysis confirmed stimulation history as the largest unique contribution (ΔR^2^ = 0.073 ± 0.056, mean±s.d.; p < 10^−15^, Wilcoxon signed-rank test; Fig. 5c). In contrast, spike history had negligible unique contribution (ΔR^2^ = –0.002 ± 0.008; p = 0.012), suggesting it did not meaningfully boost predictive power beyond the stimulation. Nonlinear terms, including stim-stim interactions (ΔR^2^ = 0.017 ± 0.028; p < 10^−6^) and stim-spike interactions (ΔR^2^ = 0.016 ± 0.018; p < 10^−11^), had small but significant contributions.

**FIG. 5.**
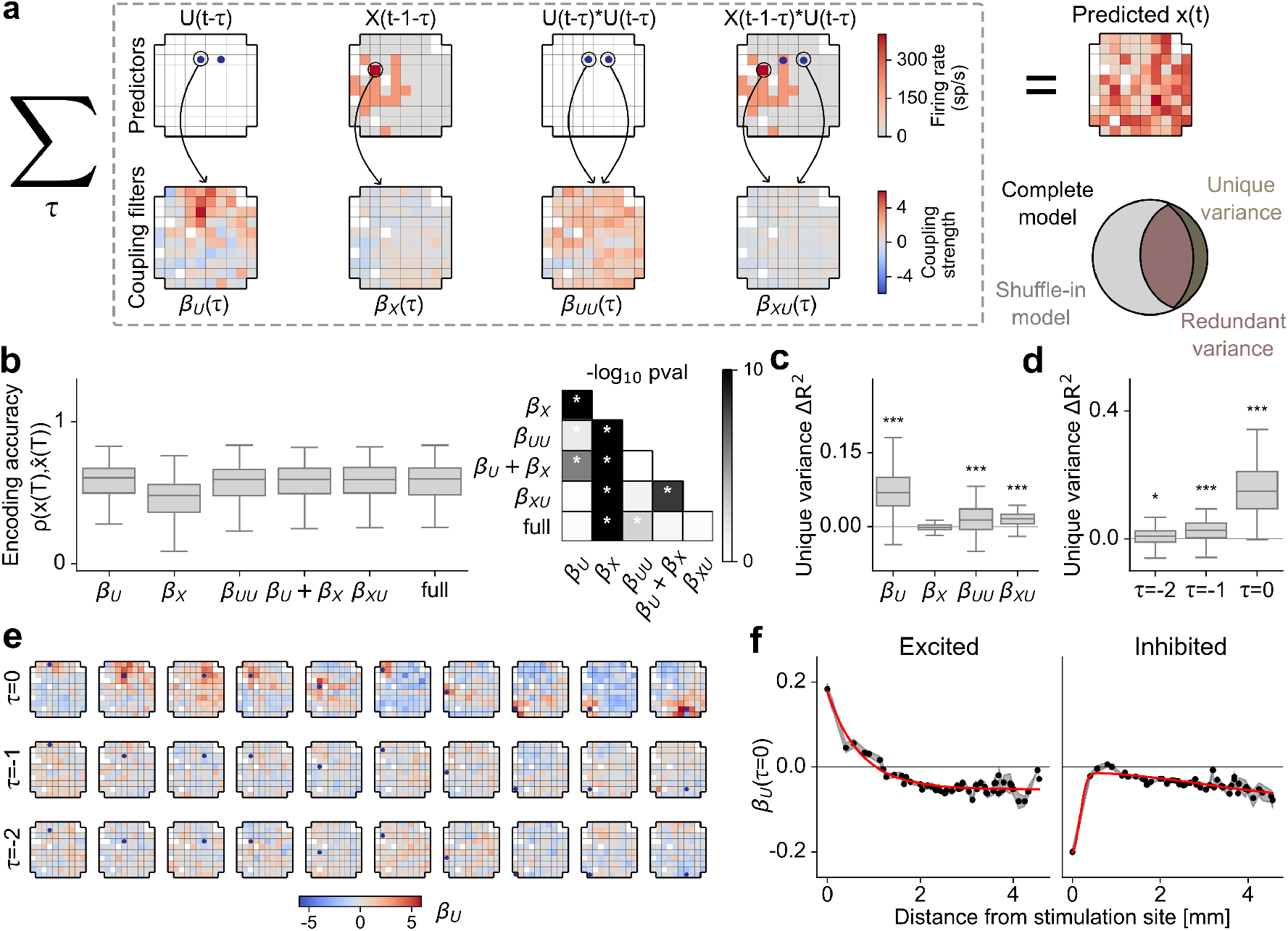
Encoding model explains micro-stimulation effects as compositions of single-electrode stimulation fields. **a)** Schematic of the linear encoding framework. Neural responses x(t) are modeled as a weighted sum of stimulation history (*β*_U_), spike history (*β*_X_), and nonlinear interactions (*β*_UU_, *β*_XU_). Unique variance for each predictor is computed by comparing the full model to a “shuffle-in” model in which the predictor is shuffled across trials. **b)** Left: cross-validated encoding accuracy (Pearson correlation) for single-feature models and the full model. Right: statistical comparisons between models. Grayscale indicates – log_10_ p-values. **c)** Unique variance (ΔR^2^) contributed by each predictor set. Stimulation history (*β*_U_) provides the largest contribution (ΔR^2^ = 0.073 ±0.056, mean± s.d.; p *<* 10^−15^). Nonlinear interactions show small but significant contributions (p *<* 10^−6^). **d)** Time-resolved unique variance for individual lags within the REACH-Ctrl forward model X = *β*_U_U. Predictive power is highest for the last stimulation (*τ* = 0, ΔR^2^ = 0.176 ± 0.139; p *<* 10^−16^) and decreases for earlier pulses (*τ* = –1, –2), with each lag providing a statistically significant and distinct contribution. *: p *<* 0.05, ***: p *<* 10^−3^, see main text for exact p-values. **e)** Representative spatiotemporal stimulation fields (*β*_U_) across three time lags (rows) for different stimulating electrodes (columns; dots show stimulating electrodes in the Utah array). **f)** Spatial profile of stimulation fields at *τ* = 0. Stimulation can excite or inhibit activity in the neural cluster recorded by the stimulating electrode. Excitatory fields exhibit exponential decay with distance from the stimulation site (left; length constant *λ* = 0.71 mm). Inhibitory fields exhibit a central suppression and a surrounding rebound (difference-of-Gaussians model with *σ*_a_ = 0.16 mm, *σ*_b_ = 2.96 mm). Gray shading indicates s.e. across electrodes.

These results indicate that the effect of stimulating multiple electrodes simultaneously is well approximated by the linear sum of the effects of stimulating each electrode alone. In matrix form, the encoding model X = *β*_U_U corresponds to the forward model of REACH-Ctrl, so the success of this linear encoding model explains why linear control theory is effective in our low-current microstimulation regime.

To further dissect the temporal dynamics within this forward model, we performed a similar estimation of unique variance for individual time steps. Predictive power was highest for the last pulse (τ = 0, ΔR^2^ = 0.176± 0.139; p < 10^−16^), and decreased for earlier pulses (τ = –1 : ΔR^2^ = 0.026 ± 0.036, p < 10^−7^; τ = –2 : ΔR^2^ = 0.007± 0.031, p = 0.026; Fig. 5d). Each lag nonetheless provided a statistically significant and distinct contribution to the overall model accuracy, consistent with a regime in which individual pulse effects partially decay and sum approximately linearly over the timescales we probed.

The encoding filters define “stimulation fields,” which describe the spatiotemporal spread of microstimulation effects across the array (Fig. 5e). We categorized these fields as “excited” or “inhibited” based on the sign of the response change at the stimulating electrode. For excited electrodes, we observed a rapid decline of excitation effects in neighboring electrodes, well characterized by an exponential decay (characteristic length λ = 0.71 mm). In contrast, inhibited electrodes exhibited a central suppression field surrounded by a rebound field, well captured by a difference-of-Gaussians model (σ_a_ = 0.16 mm, σ_b_ = 2.96 mm). These localized spatiotemporal dynamics are consistent with the characteristic length scales of stimulation effects reported in prior studies [43] and demonstrate that microstimulation recruits distinct spatial patterns of activation and suppression depending on the local neural context.

## Discussion

Developing model-based strategies to control population activity patterns is a central challenge in systems neuroscience. Here we show that a closed-loop brain–computer interface can achieve real-time control of spiking activity in macaque prefrontal cortex using only short training epochs of microstimulation and population recordings. Our REACHable manifold Control algorithm (REACH-Ctrl) learns, from random multi-electrode pulse sequences and their evoked responses, which population states are reachable and which stimulation patterns steer the circuit toward designated targets. This data-driven framework bypasses the need for an explicit model of connectivity or dynamics, yet delivers high control accuracy using only single-session data. By working directly with input-output maps derived from clinically relevant microstimulation hardware, REACH-Ctrl links control theory to cortical population activity in a way that is both experimentally practical and directly compatible with translational neuromodulation.

Most quantitative approaches to network control assume knowledge of a dynamical model or structural connectome [19–22, 26]. In intact cortex, this information is not available in vivo: inferring a high-dimensional recurrent network from limited, noisy recordings is a non-identifiable inverse problem, with many distinct dynamical systems reproducing the same trajectories under sparse sampling [24, 25]. REACH-Ctrl bypasses this bottleneck by working directly with input–output data. Using recent results in data-driven control [38–41], it constructs a controllability map from random multi-electrode stimulation sequences to their evoked responses, then uses this map to design low-current stimulation patterns that steer the population along desired trajectories.

A striking feature of our results is that this linear, data-driven framework performs well in cortical circuits that are, in general, nonlinear and state-dependent. Our encoding analyses clarify why. In the weak microstimulation regime we adopt, population responses to multi-electrode sequences are well approximated by the linear sum of single-electrode “stimulation fields,” with only modest contributions from nonlinear interactions and spike history. The linear component captures most of the explainable variance, and the same linear map underlies the controllability matrix used by REACH-Ctrl. Together with modeling work on linearization around operating points [23, 53], this suggests that under low currents, short horizons, and a modest number of simultaneously stimulated electrodes, cortical dynamics are effectively linear from the perspective of the stimulation–response mapping we exploit. Outside this regime—at higher currents, longer horizons, or with more densely coordinated pulses—nonlinear phenomena such as synaptic saturation, state transitions, and oscillations are likely to emerge and would require extensions of the present framework.

The manifold analyses clarify what is being controlled. The intrinsic manifold, defined by pre-stimulation activity, captures the dominant modes of variability expressed during spontaneous activity [54–56]. The reachable manifold, defined by the data-driven controllability matrix, captures the repertoire of patterns that can be produced by microstimulation. We find substantial overlap between these manifolds, but also systematic differences: multi-pulse sequences generate trajectories with components both within the shared subspace and in directions that lie in the portion of the reachable manifold orthogonal to the intrinsic manifold. Thus, REACH-Ctrl can both exploit intrinsic modes and drive the circuit into off-manifold states [57], in contrast to many volitional BCI paradigms in which activity remains tightly constrained to intrinsic manifolds [58]. We developed a new visualization method, Gramian Component Analysis (GCA), that formalizes this distinction by ranking axes by how easily they can be reached, rather than by how much variance they explain.

Our choice of electrical microstimulation is driven by translational considerations. Optical and optogenetic methods, including two-photon imaging and holographic photostimulation, provide exquisite spatial and cell-type specificity in rodents and other small animals [13–15, 28, 29, 33], but they require optical access and genetic tools that are not yet available for routine use in humans and face substantial practical and ethical barriers. Electrical microstimulation and deep brain stimulation lack single-cell specificity and engage mixed populations via complex combinations of local and long-range effects [35, 36, 46], yet they are already widely deployed for movement disorders, epilepsy, and refractory psychiatric conditions [1, 4–9]. By showing that sophisticated, model-based control can be implemented with microstimulation through standard Utah arrays or comparable devices, our work helps bridge the gap between mechanistic circuit experiments and clinically relevant neuromodulation. Crucially, our approach is extremely data-efficient as it leads to successful control leveraging single-session training data only (for approaches based on amortization see [59]).

Our results establish a proof of principle: precise, sample-efficient control of cortical population activity is feasible with clinically relevant hardware and realistic recording constraints, without an explicit model of circuit dynamics. The present implementation is limited to short horizons, weak perturbations, and resting states; repeated stimulation, neuromodulatory context, and behavior are likely to induce plasticity and state dependence that will require adaptive or state-dependent controllers. Extending REACH-Ctrl to task-engaged states, and ultimately to shaping behavior, is a key next step for realizing the full translational potential of this framework. Nonetheless, by combining data-driven controllability, a geometric description of intrinsic and reachable manifolds, and empirical characterization of stimulation fields, our framework provides a general blueprint for designing perturbations in sparsely observed neural circuits *in vivo*. Because REACH-Ctrl operates with stimulation hardware already used in humans, it also offers a concrete path toward model-based neuromodulation strategies that are informed by, rather than blind to, the population dynamics they seek to control.

## Methods

We recorded from and electrically stimulated the prearcuate gyrus (area 8Ar, Fig. 1) of two macaque monkeys (*Macaca mulatta*) during quiet wakefulness. All experimental procedures were approved by the Institutional Animal Care and Use Committee at New York University and conformed to the National Institutes of Health *Guide for the Care and Use of Laboratory Animals*.

During experiments, monkeys were seated in a semi-dark room with their heads stabilized using a surgically implanted head-post. Recording and stimulation were performed through a chronically-implanted Utah array (96 electrodes; Blackrock Neurotech) [60–62]. Array electrodes were 1 mm long with 400 µm inter-electrode spacing, permitting simultaneous recordings from neighboring cortical microcolumns in a 4 × 4 mm^2^ region of cortex (Fig. 1). Voltage signals recorded by the electrodes were bandpass filtered and thresholded online with Blackrock’s Cerebus recording system to identify neuronal spikes. Spike waveforms as well as raw voltage signals were saved for offline processing (sampling rate, 30 kHz). Eye position was monitored at 1 kHz using a high-speed infrared eye-tracking system (Eyelink, SR-Research, Ontario) to confirm that monkeys remained awake throughout the experiments.

Electrical microstimulation was delivered through the Utah array electrodes using a CereStim stimulator (Blackrock Neurotech). Microstimulation pulses were low-current (15 µA) and biphasic with a total duration of 200 µs (Fig. 1). Our experimental setup allowed delivery of trains of microstimulation pulses through individual electrodes or arbitrary combinations of electrodes. At each pulse, we could simultaneously stimulate any combination of up to 16 electrodes, generating diverse spatial patterns. Different patterns were delivered at predefined temporal intervals (12.5-50 ms), while the system rapidly switched between stimulation and recording modes to collect spiking activity between stimulation pulses (switching latency, 3-5 ms). The number of electrodes stimulated in each pulse and the number of pulses delivered in each trial were determined by REACH-Ctrl (see below).

Microstimulation of the prearcuate gyrus with currents >50 µA can trigger saccadic eye movements [44]. We used low-current microstimulation, well below this motor threshold, to avoid stimulation-induced saccades. Consistent with this, we never observed saccades time-locked to microstimulation in our experiments. The low stimulation current also limited the number of affected neurons, improving the spatial specificity of stimulation effects and minimizing the risk of electrode damage with repeated stimulations.

Because Utah arrays are already in widespread use in human intra-cortical studies [37, 63–67], our methods are compatible with translation to human applications.

A typical session consisted of three blocks: a resting-state block in which we recorded neural activity to identify electrodes with the largest stimulation effects on the circuit [43] (duration, 12-20 min), a training block in which we recorded the effect of random stimulation patterns on the circuit activity (duration <5 min), and a test block in which we delivered a set of stimulation sequences to compare the evoked and target activity patterns (duration, 15-30 min). REACH-Ctrl used the training data to determine the reachable manifold and to generate target stimulation sequences (see below). The training and test blocks were divided into trials. Trial duration was fixed within a session but could vary across sessions (range, 1.7-7 s). The stimulation sequence was delivered at a random time within each trial. Stimulation onsets on consecutive trials were separated by at least 1.5 s to give the circuit ample time to settle following each stimulation.

## Removal of stimulation artifacts

Switching between recording and stimulation modes caused brief electrical artifacts that settled within milliseconds. To remove these artifacts and ensure they did not influence our analyses, we developed an data-driven artifact detection algorithm. For the training and test blocks, we defined three sets of non-overlapping time windows: reference windows spanning 500 ms before stimulation sequence onsets, validation windows from 200 to 700 ms after the last pulse in the sequence, and analysis windows between consecutive pulses. The reference and validation windows were artifact free and provided a dictionary of valid spike waveforms. For each electrode, we first characterized the variability of artifact-free waveforms. We computed the distance between every spike waveform in the validation windows and all spike waveforms in the reference windows. In this reference-validation distance matrix, each column contained the distances from a validation spike to all reference spikes, providing an empirical embedding of artifact-free waveforms. We then embedded waveforms recorded in the analysis windows in the same space by computing their distances to all reference spikes. For each analysis-window waveform, we compared its distance to the nearest validation waveform in the embedding space with the distribution of nearest-neighbor distances among validation spikes themselves. Waveforms whose nearest-neighbor distance exceeded this distribution by a preset threshold (see below) were rejected as putative artifacts.

We use two distance metrics: Euclidean distance between spike waveforms and their correlation distance 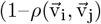, where 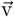 is the vector of spike waveform voltages sampled at 30 kHz). Analysis window waveforms whose distance to the nearest validation spike waveform exceeded the threshold for either metric were excluded. The threshold for the correlation distance was set to the 99.5th percentile of nearest-neighbor distances among validation spikes. The threshold for the Euclidean distance was set to the 99.95th percentile. These thresholds were chosen to reject all artifacts while minimizing the exclusion of valid spike waveforms. An experienced electrophysiologist (I.M. or R.K.) visually confirmed that rejected waveforms corresponded to stimulation artifacts.

Artifacts removal was performed automatically after each task block to ensure REACH-Ctrl was not affected by stimulation artifacts. Firing rates in the inter-pulse intervals were quantified in windows were valid spikes dominated: from 7.5 ms after the stimulation pulse until the next pulse onset.

### Identification of high-efficacy electrodes

We began each session with a resting epoch prior to the training block, during which we recorded spontaneous spiking activity while the monkey sat quietly with eyes open. Immediately after, we applied the Causal Flow (CF) algorithm [43] to infer directed functional connectivity among electrodes (Supplementary Fig. S1a). CF revealed a hierarchical structure among the neural clusters recorded by different electrodes: a small subset of electrodes (“hubs”) exhibited markedly elevated directed functional connectivity to other electrodes (Supplementary Fig. S1b). The selection of stimulated electrodes was based on the hub ranking determined by the CF algorithm which was run at the end of the resting epoch and preceding the training epoch (approximate runtime 10 min). In separate experimental sessions (n=17), we confirmed that stimulation of hub electrodes produced stronger activity modulations across the array (Supplementary Fig. S1c,d), consistent with past studies [43, 68]. Moreover, CF hubness estimation was consistent across months in both monkeys (Supplementary Fig. S1e)

### REACH-Ctrl

#### Overview and training data

To evoke a target ensemble activity vector at a fixed horizon T using a T-step microstimulation sequence, we developed REACH-Ctrl. REACH-Ctrl builds on data-driven control methods that enable controller design directly from input-output data, without requiring an explicit dynamical model or connectivity [39–41]. In each session, we first collected a training dataset of stimulation-response pairs. Specifically, during the training block we delivered N_train_ random multi-electrode stimulation sequences U = [u^(1)^, …, u^(Ntrain)^], where each u^(k)^ ∈ {0, 1}^mT^ is an mT-dimensional binary vector indicating which of the m electrodes were stimulated at each of the T steps on trial k (with u^(k)^ reshaped into T consecutive blocks of length m, one block per pulse). For each trial k, we recorded the corresponding population responses 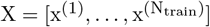, where x^(k)^ ∈ℝ^nT^ is the concatenated n-dimensional population spike-count response across the T steps of trial k.

Responses were defined as population spike counts in 6 ms bins after each microstimulation pulse, beginning 5 ms after pulse onset. From X we extracted the final-step responses 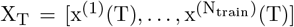, where x^(k)^(T) ∈ ℝ^n^ denotes population spike-count vector following the last pulse of the T-pulse sequence on trial k. Each pair (u^(k)^, x^(k)^(T)) provides an input-output example linking a length-T stimulation sequence to the resulting population state at horizon T.

#### Controllability matrix estimation

The fundamental object in REACH-Ctrl is an n *×* mT data-driven controllability matrix C_T_ that maps any T-step input sequence u to the population activity state at time T,

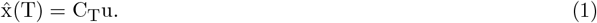

An unregularized estimate of C_T_ can be obtained directly from the training data by least squares,

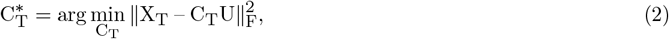

whose solution is 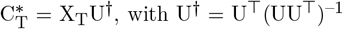, with U^*†*^ = U^*T*^(UU^*T*^)^−1^ the Moore-Penrose pseudoinverse when U has full row rank. For identifiability, the stimulation design matrix U must have full row rank (i.e., rank(U) = mT), which is satisfied with high probability for random i.i.d. training sequences when N_train_ ≥mT [40, 41] (“rank condition”). In the Supplementary Text, we detail the assumptions under which this data-driven formulation recovers the reachable subspace and connects to optimal control for linear time-invariant systems.

In the presence of noise, the unregularized estimate can have high variance. We therefore used a ridge-regularized estimator,

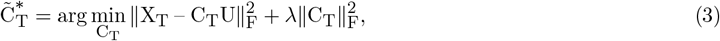

with closed-form solution 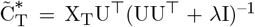. We selected λ by five-fold cross-validation on the training dataset. The resulting 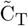 was used online and improved held-out control accuracy (Supplementary Fig. S2a).

#### Reachable manifold and target selection

The column space of 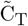 determines the set of final states that can be produced (in the linear model) by length-T stimulation sequences. To define a low-dimensional reachable manifold and prioritize directions that are most strongly driven by stimulation, we computed the left singular vectors of 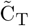 (equivalently, the eigenvectors 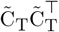) and retained directions associated with nonzero singular values. We then chose target ensemble activity patterns x_f_ as linear combinations of the top singular vectors, corresponding to directions that are most easily reachable under the learned mapping.

#### Optimal controller design (unconstrained)

Controller design follows data-driven control results based on Willems’ Fundamental Lemma [47]. Under the assumptions explained in the Supplementary Text, any reachable final state at horizon T can be represented from the training data by choosing coefficients α such that x_f_ = X_T_α, which induces the corresponding input u = Uα. The minimum-energy controller in this parameterization is obtained by solving

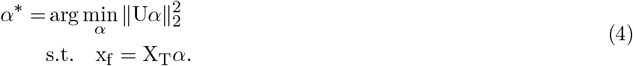

Equivalently, one can write the optimization directly over u:

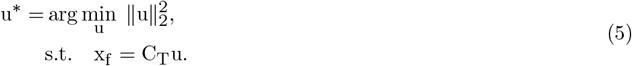

#### Input constraints and binary optimal controllers

In our experimental setting, inputs were constrained by the stimulation hardware and safety considerations. Each electrode was either stimulated at a fixed current amplitude or not stimulated, motivating a binary input representation. In addition, to limit injected current we constrained the number Q of simultaneously stimulated electrodes per time step (typically Q = 2; in some sessions we allowed Q ∈{1, 2, 3}). Operationally, we reshaped u ∈ ℝ^mT^ into T blocks u_t_ ∈ℝ^m^ and enforced sparsity within each block.

We implemented two approaches to obtain constrained controllers. In the first approach, we computed the standard minimum-energy solution using the regularized estimate,

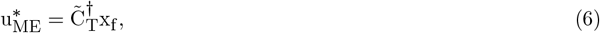

and then discretized it by selecting, within each time step t, the Q largest entries of u^*ME^ (after reshaping into T blocks) and setting all other entries to zero.

In the second approach, we incorporated quantization and sparsity constraints more directly via a relaxed nonconvex optimization. We minimized a binarization penalty while encouraging accurate target matching and enforcing the desired total sparsity, using the box constraint 0 ≤ u ≤ 1:

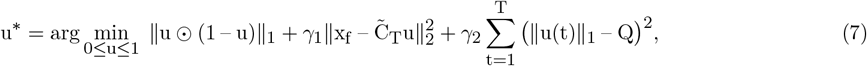

where ⊙ denotes the element-wise (Hadamard) product and γ_1_, γ_2_ control the tradeoff between target matching and input constraints. Because quantization generally makes x_f_ = 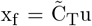 unattainable, we defined the effective target for each computed controller as the model-predicted final state

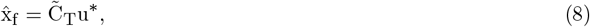

and evaluated control performance relative to 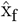 (unless otherwise stated). Since the relaxed optimization is nonconvex, to mitigate local minima we initialized from 100 random starting points and retained the solution whose predicted final state 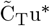 was closest to the desired x_f_.

In each session, the full REACH-Ctrl procedure—estimation of 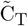, selection of target patterns, and computation of a constrained controller for each target—was run online at the end of the training epoch (runtime ∼30 s). Both controller-construction approaches yielded similar control accuracy (Supplementary Fig. S2).

### Control accuracy

To quantify the accuracy of REACH-Ctrl, we compared model predictions with evoked activity in the test block. For each stimulation sequence, we computed the predicted target population activity as 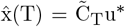, where u^*^ is the optimal input sequence and 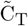 is the regularized controllability matrix in Eq. 3, estimated from the random stimulation-response data of the training block. Control accuracy for each sequence was defined as the Pearson correlation 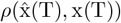 between the predicted activity 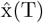 and the evoked population activity x(T), defined as trial-averaged spike counts across channels for each test sequence.

To establish significance, we compared the predictive accuracy to a null distribution obtained from surrogate datasets where the controllability matrix was derived by randomly permuting training labels X_T_. This procedure disrupts the alignment between stimulation and evoked responses while preserving the stimulation and response statistics. The shuffling process was repeated N_shuffle_ = 10^3^ times to obtain the null distribution. At the session level, the p-values were computed as the fraction of shuffled accuracy exceeding the empirical accuracy. To assess the statistical significance across all sessions, we performed a Wilcoxon signed-rank test comparing the observed experimental accuracies against the mean of the shuffled distributions for each session. Since experimental results were qualitatively consistent across both monkeys, data were pooled.

### Gramian Component Analysis

To visualize neural trajectories, we developed Gramian Component Analysis (GCA), a new dimensionality reduction method tailored to control. GCA starts from the data-driven controllability matrix 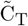 and computes its left singular vectors, ordered by decreasing singular values. These singular vectors identify directions in state space that can be most efficiently driven by stimulation. Unlike principal component analysis (PCA), which reflects only the variance of observed activity, GCA explicitly incorporates the stimulation-response mapping encoded by 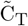, and thus projects activity onto the “reachable manifold,” the subspace most relevant for control (see Supplemental Information for the relationship between GCA and PCA).

### Pattern discriminability

To quantify whether evoked responses in the test epoch were sequence-specific, we performed a cross-validated classification using linear discriminant analysis (LDA). For each trial, we used spike counts across electrodes at each time step (including pre-stimulus activity at t = 0) as features, and tretaed combinations of stimulation sequence and time step as labels (Fig. S3). In Fig. 3, spike counts at the last time step were used as features and stimulation sequences as labels. LDA performance was evaluated with 5-fold cross-validation on test-epoch trials. Statistical significance was assessed by comparing the classification accuracy in the empirical data to a null distribution obtained from surrogate datasets generated by shuffling labels across trials (N_shuffle_ = 10^3^).

### Fraction of reachable-manifold energy explained by neural trajectories

#### Reachable manifold energy and its projection onto directions spanned by data

Let ℳ_A_ ⊂ℝ^m^ denote the linear subspace (“manifold”) spanned by the columns of the matrix A ∈ℝ^m*×*n^. We interpret the matrix AA^*T*^ as defining a *covariance* (or second-moment) operator on activity space, with total variance

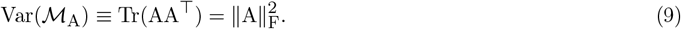

Given a unit vector v ℝ^m^, the quantity v^*T*^AA^*T*^v measures the variance of ℳ_A_ along direction v. We quantify how much of the manifold variance lies in directions spanned by observed neural trajectories by summing this directional variance over an orthonormal basis for the trajectory subspace.

#### Variance explained by a single direction

For a nonzero vector x ∈ ℝ^m^, define its associated unit direction v = x/ ∥x∥_2_. The fraction of manifold variance aligned with direction x is

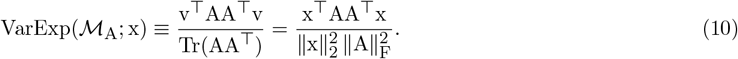

This quantity is invariant to rescaling of x and satisfies 0 *≤* VarExp(ℳ_A_; x) *≤* 1.

#### Variance explained by a neural trajectory

Let X = [x_1_ *…* x_t_] ∈ ℝ^m*×*t^ be a collection of (possibly collinear) vectors, and let *S*_X_ = span(X) denote the subspace they span. Let Q_X_ ∈ ℝ^m*×*q^ be an orthonormal basis for *S*_X_ (obtained by SVD; see below, where we choose q by variance retention). The fraction of manifold variance explained by *S*_X_ is defined as the total variance of ℳ_A_ within *S*_X_ normalized by the total variance of ℳ_A_:

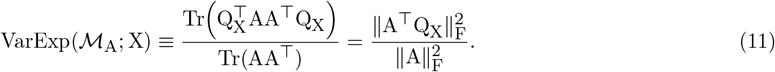

If *S*_X_ contains all left singular vectors of A (i.e., ℳ_A_ *⊆ S*_X_), then VarExp (ℳ_A_; X) = 1. More generally, VarExp(ℳ_A_; X) measures how much of the energy of ℳ_A_ lies in directions spanned by X.

#### Controlling for collinearity in X

To avoid overweighting redundant (collinear) columns of X, we compute the thin SVD X = UΣV^*T*^ and retain the smallest q such that

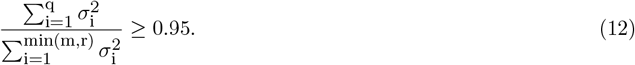

where r = rank(X) and σ_i_ are the singular values. We then set Q_X_ = U_q_, where U_q_ ∈ ℝ^m*×*q^ retains the first q columns of U, and evaluate Eq. 11 as

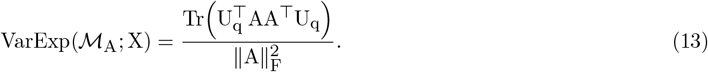

#### Variance within shared vs. non-shared components of two manifolds

Let ℳ_A_ and ℳ_B_ have orthonormal bases 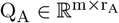 and 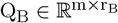 (e.g., left singular vectors of A and B with nonzero singular values). Principal angles are obtained from

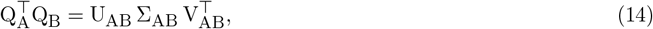

where diagonal entries σ_AB,i_ are the cosines of principal angles. We defined “shared” directions as those with principal angle at most 15^*°*^, i.e. σ_AB,i_ *≥*τ with τ = cos(15^*°*^) ≈0.966. Let I = {i : σ_AB,i_≥ τ} and define an orthonormal basis for the shared subspace as

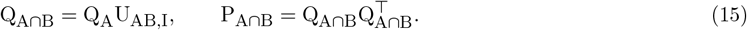

The component of ℳ_A_ not shared with ℳ_B_ is defined by

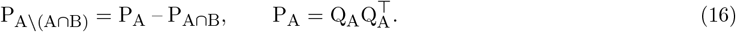

We then quantify the fraction of ℳ_A_ variance that lies in its shared vs. non-shared components (optionally restricted to the directions spanned by X). Specifically, the fraction of ℳ_A_ variance contained in the shared subspace is

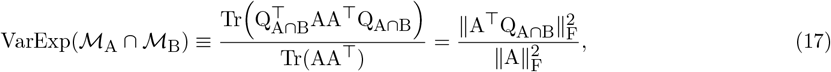

and an analogous expression holds for the non-shared component by replacing Q_A∩B_ with any orthonormal basis of ℳ_A \_ (ℳ_A_ ∩ℳ_B_).

To quantify the fraction of ℳ_A_ variance that lies in the shared component *and* is expressed along the directions spanned by X, we combine both restrictions by projecting the trajectory subspace onto the shared basis:

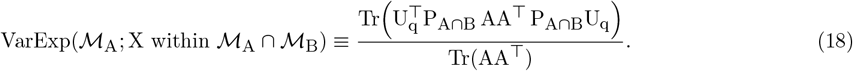

An analogous expression holds for the non-shared component by replacing P_A∩B_ with P_A\ (A∩B)_.

#### Application to reachable and intrinsic manifolds

We applied these definitions to 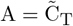 (reachable manifold) and 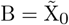 (intrinsic manifold), where X_0_ ∈ℝ^neurons×trials^ is the pre-stimulus spike-count matrix (5 ms bin immediately before the first pulse) and 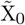 is its reduced-rank approximation. For each session and each time t, we formed he matrix of trial-averaged evoked responses in the test epoch by concatenating evoked activity across stimulation sequences and time points,

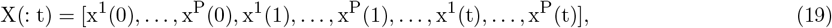

where P is the number of stimulation sequence patterns in the session. We computed VarExp(ℳ_reach_; X(: t)) and its shared and non-shared decompositions using the equations above.

### Encoding models

To gain a mechanistic understanding of how data-driven control shapes neural activity, we fit encoding models to predict population responses from stimulation history and recent spiking history. Such encoding models have been shown to capture population responses to visual stimuli, task variables, and spike history [49–52, 69]. Inclusion of stimulation history U and spike history X as predictors enabled our models to capture both the effects of external stimulation and recurrent temporal dependencies between electrodes. To account for nonlinear effects, the design matrix also included pairwise interactions between between stimulated electrodes and between stimulation and spike history. These terms allow the model to capture nonlinear dependencies, such as how the effect of stimulating one electrode may depend on the stimulation or spiking activity state of other electrodes.

The full encoding model is expressed as

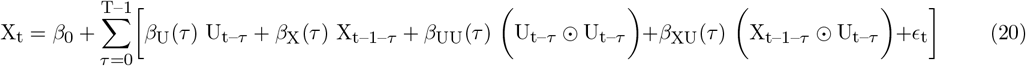

where X_t_ and U_t_ encode the population activity and stimulation pattern at time t across all training trials, *β*_k_ are model coefficients, ⊙denotes element-wise products, ϵ_t_ is the residual, and T is the number of steps in the microstimulation pulse sequence (T = 3 in most sessions). The model uses only past and current predictors, respecting causality.

The model was fit to the single-trial data collected in the training block using ridge regularization, with the regularization parameter selected via five-fold cross-validation. Alternative regularization methods yielded comparable prediction accuracy (Supplementary Fig. S5).

Within the full encoding model, we quantified both the overall and unique contributions of each set of predictors [51, 52]. To estimate the total explanatory power of each predictor set (e.g., stimulation history, spike history, interaction terms), we fit a set of reduced models that included only that predictor to the training epoch data and computed cross-validated explained variance (cvR^2^) on trial-averaged population activity from the test block. This measure captures how much of the mean response pattern each predictor set can explain on its own, providing an upper bound on the predictive power of the predictor set. However, because of potential co-linearity between different predictor sets, cvR^2^ is not, by itself, a measure of unique contribution.

To quantify the unique contribution of each predictor set, we followed a shuffle-in-model approach. For each set of predictors, we shuffled that predictor across trials within the full model while leaving all other predictors intact (N_shuffle_ = 10^3^). The reduction in explained variance relative to the full model, ΔR^2^, captures the unique variance explained by that predictor set.

We also examined the temporal structure of stimulation effects by applying the same procedure to individual lags τ within the stimulation predictors, quantifying cvR^2^ and ΔR^2^ for individual stimulation pulses. This analysis identified which pulses in the sequence were most informative for the population activity pattern at the end of the sequence.

To ensure robustness of our findings, we compared the linear encoding model to alternative model variants, including Poisson GLMs and reduced-rank regression (Supplementary Fig. S5). Poisson GLMs account for the discrete and non-Gaussian nature of spiking activity, and reduced-rank regression constrains solutions to a shared latent space. These variants did not improve predictive performance and yielded similar patterns of feature importance, cvR^2^, and ΔR^2^. In the Supplementary Text, we show that the Poisson GLM reduces to the linear encoding model in Eq. (20) in the regime of parameters and observations used in our experiments.

## Acknowledgments

This work was supported by the National Institute of Mental Health (R01 MH127375 and R01 MH141929). R.K. and I.M. were additionally supported by the Pew Innovation Fund (00037214).

## Author Contributions

G.B., A.D., I.M., L.M., and R.K. conceptualized and designed the study. I.M. collected the data. G.B., A.D., and C.C. analyzed the data. G.B., A.D., L.M., and R.K. wrote the manuscript.

## Data availability

All datasets generated and analyzed in this study will be deposited in a public repository and made available upon publication.

## Code availability

All analysis code will be deposited on Github and made available upon publication.

## Supplemental Text

### A Gramian control of linear time-invariant systems

To formalize how patterned microstimulation shapes population spiking activity, we modeled the neural population dynamics as a discrete-time, linear dynamical system:

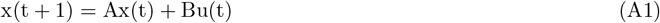

where x(t) is an n-dimensional state vector describing the population spike counts at time t, u(t) is an m-dimensional vector encoding microstimulation on a subset of m electrodes, A is the n × n intrinsic coupling matrix between neural clusters recorded by the electrodes, and B is the n × m input matrix specifying how stimulation drives the system.

In the main manuscript we consider systems with vanishing initial conditions x_0_ = 0. In this case, the state at time can be written as x(T) = C_T_u, where

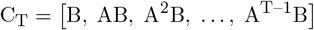

is the finite-horizon controllability matrix, and u stacks the input vectors over time. We define the *reachable manifold* as the set of states reachable at time T, given by ℛ_T_ = Im(C_T_) where Im(C_T_) denotes the column space of C_T_. A target state x_f_ is reachable in T steps if there exists an input sequence u such that x_f_ = C_T_u.

Minimum-energy control seeks an input sequence u* = [u_0_, …, u_T–1_] that steers the system toward x_f_ in T time steps while minimizing the control energy 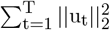. For a controllable system, the minimum-energy control sequence is

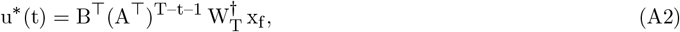

where the finite-horizon controllability Gramian

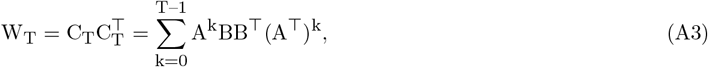

quantifies how easily different directions in state space can be driven.

Although these closed-form solutions for the optimal controller and Gramian are theoretically appealing, they can be rarely used directly in practice because they require accurate knowledge of A and B, they are sensitive to model uncertainty, and the controllability Gramian is often ill-conditioned in realistic high-dimensional systems [40, 70, 71]— the optimal controller becomes numerically unstable as it requires inverting large matrices with potentially small eigenvalues. These limitations motivate a data-driven approach that bypasses explicit estimation of A and B.

### B Data-driven control

A data-driven alternative to Gramian-based control exploits input–output measurements to reconstruct finite-horizon controllability properties without explicitly identifying the system matrices. Consider a collection of N trials in which random input sequences u^(k)^ are delivered in trial k and the state at time T is observed

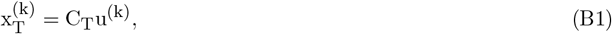

We collect stimulation inputs and terminal activity states into data matrices U = [u^(1)^, …, u^(N)^] and X_T_ = [x^(1)^, …, x^(N)^]. Building on Willems’ Fundamental Lemma, we assume that the optimal input can be computed as a linear combination of the columns u^(k)^ of U. Within this representation, the minimum-energy control problem admits a purely data-driven solution. If U has full row rank, i.e.,

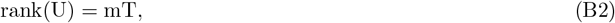

then the optimal input sequence steering the system to a desired target x_f_ in T steps is given by [40]

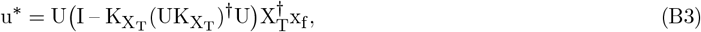

where 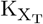 spans Ker(X_T_). This expression is equivalent to

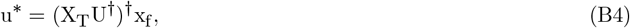

from which we can define the finite-horizon data-driven version of the controllability matrix as

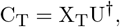

and the corresponding data-driven controllability Gramian,

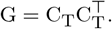

When U is not full row rank, the minimum-energy solution remains well defined provided u^*^ ∈ Im(U). Otherwise, the controller drives the system to the closest reachable state 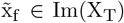 Im(X_T_), reflecting the fact that reachability is constrained by the subspace spanned by the experimental data.

### C Equivalence between linear and nonlinear encoding models in the limit of small amplitude stimulation

While REACH-Ctrl utilizes a data-driven linear framework, neural population dynamics are fundamentally nonlinear, and are often modeled as Poisson processes where firing rates are governed by non-negative activation functions. To justify the linear approximation, we consider a general nonlinear encoding model where the population state x(T) at time T is a nonlinear function of the accumulated stimulation:

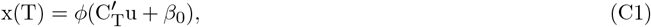

where *ϕ*(·) is a monotonic, non-negative activation function (e.g., exponential or sigmoid), and *β*_0_ represents the baseline drive (encapsulating the initial state and constant offsets). In the limit of small stimulation amplitudes (u → 0), a first-order Taylor expansion allows the response to be approximated as:

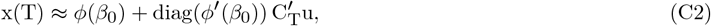

where ϕ^′^(*β*_0_) is the vector of element-wise derivatives evaluated at *β*_0_, and diag(ϕ^′^(*β*_0_)) denotes the diagonal matrix with entries 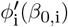.

This formulation is equivalent to a linear model with controllability matrix 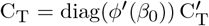. Crucially, this shows that the parameters governing the nonlinear response are simply a rescaling of the data-driven controllability matrix C_T_ by the local gain of the activation function, ϕ^′^(*β*_0_). This derivation demonstrates that for small-amplitude inputs, the nonlinear encoding model is locally equivalent to the linear case. This equivalence justifies the use of a linear control framework as a robust and computationally efficient approximation for microstimulation-driven neural control.

### D Relation between GCA and PCA

Gramian Component Analysis (GCA) and Principal Component Analysis (PCA) both provide low-dimensional embeddings of high-dimensional neural activity, but they emphasize fundamentally different properties of the data. PCA identifies directions of maximal variance by diagonalizing the empirical covariance matrix and capturing the dominant direction of variance in the population activity fluctuations, regardless of whether these fluctuations are influenced by stimulation. In contrast, GCA is grounded in control theory and focuses specifically on the directions of state space that can be efficiently driven by external inputs. In the data-driven formulation, the controllability Gramian is estimated as

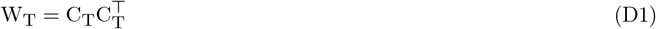

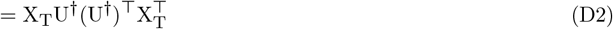

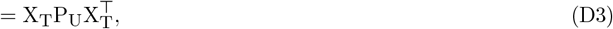

where the matrix P_U_ = U^†^(U^†^)^⊤^ acts as a weighting kernel derived from the input patterns explored during the experiment. Thus, rather than using the full covariance 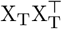, GCA computes a *projected covariance* that retains only the component of the neural variability that is explainable, and therefore controllable, by the delivered stimulation patterns. This distinction is especially important in our microstimulation experiments, where inputs act through a small number of electrodes or channels. In such settings, PCA may be dominated by uncontrolled or spontaneous variability, whereas GCA highlights the dimensions of neural activity that can actually be influenced by the stimulation.

The two methods become identical in the special case where the inputs are i.i.d. white noise with full rank, such that

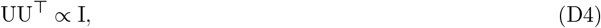

In this case, W_T_ becomes proportional to 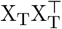 restricted to the driven subspace, and the leading GCA components converge to the leading principal components of the evoked activity. More generally, GCA can be viewed as PCA applied to a covariance matrix that has been reweighted by the structure of the inputs, so that axes are ranked by how easily they can be driven by stimulation rather than by how much they fluctuate.

## Supplemental Figures

**FIG. S1.**
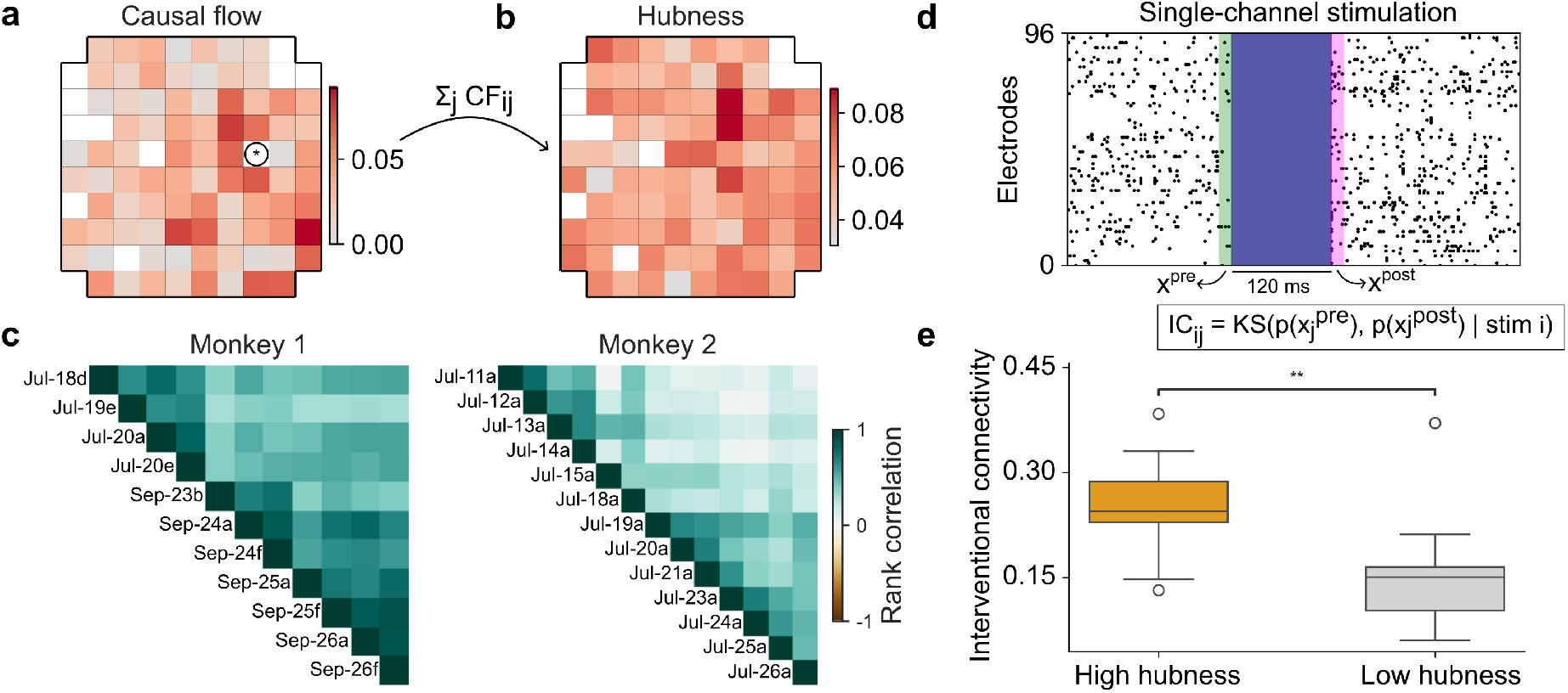
Data-driven identification of high-efficacy electrodes. **a)** Representative causal flow (CF) matrix [43] inferred from spontaneous spiking activity during the resting epoch. Each entry (i, j) represents the predicted directed functional influence from source electrode i to target electrode j. The panel shows CF in an example session for the source electrode marked by a circled star. **b)** Spatial distribution of electrode “hubness” across the 96-channel Utah array. Hubness is defined as the average CF value across target electrodes for each source electrode [43]. Hubness represents the relative strength of directed influence exerted by each electrode on the rest of the population. **c)** Temporal stability of the network structure for Monkey 1 and Monkey 2. Each heatmap shows the rank correlation of the CF-derived hubness across different experimental days. High correlation values demonstrate hubness stability over multiple weeks/months of recording. **d)** Estimation of interventional connectivity from single-channel stimulation. In n = 17 additional sessions, we perform single-channel stimulation for 120 ms (15 *µ*A biphasic pulses delivered at 200 Hz). The panel shows spiking activity before and after stimulation in a representative trial. In each session, two specific channels were selected for stimulation: one identified as a network hub (“High hubness”) and the other as a non-hub (“Low hubness”). For each source electrode i and target electrode j, we estimated pairwise interventional connectivity IC_ij_ as the Kolmogorov-Smirnov distance between the distribution of spike counts of target electrode j before (green) vs. after (magenta) stimulation of source electrode i (spike counts binned in a window of 5 ms). We further define the IC of a source electrode, IC(i), as the column average of IC_ij_ across all targets j. **e)** CF predicts the strength of stimulation effects. The IC of electrodes classified as network hubs is significantly higher than the IC of electrodes classified as non-hubs. Wilcoxon signed-rank test, p = 0.004.

**FIG. S2.**
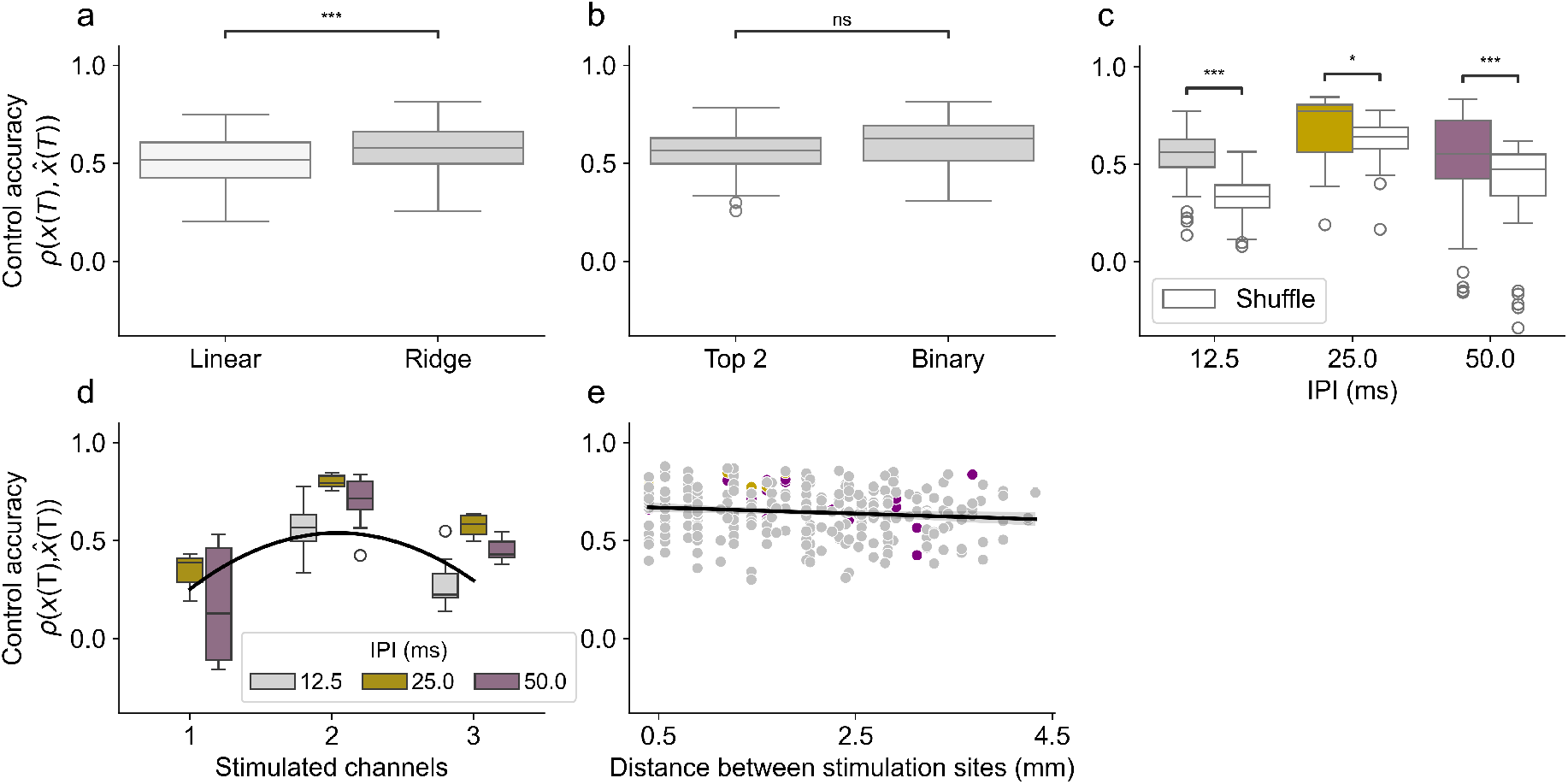
Effect of experimental parameters on control accuracy. **a)** Optimization of the controllability matrix via ridge regularization. Comparison of control accuracy between a standard linear least-squares estimation and the ridge-regularized controllability matrix. Introducing the regularization parameter λ, optimized via cross-validation, significantly improved robustness to noise and control accuracy (*ρ*_linear_ = 0.52 ± 0.13, mean ± s.d.; *ρ*_ridge_ = 0.57 ± 0.12, p < 0.001, Wilcoxon signed-rank test). **b)** Performance comparison of input-constraint strategies. Control accuracy is shown for the two approaches used to satisfy binary and amplitude constraints for our microstimulation pulse sequences: the binarized minimum-energy solution 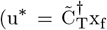 followed by truncation) and the non-convex optimization incorporating soft penalties for binary structure and total electrode count. Both methods yielded similar control accuracy (Wilcoxon signed-rank test, p = 0.61), indicating that REACH-Ctrl is robust across different numerical solvers. **c)** Control accuracy as a function of the inter-pulse interval (IPI), compared to randomized null distributions. Colored boxplots represent empirical accuracy (12.5 ms: 0.63 ± 0.13; 25.0 ms: 0.67 ± 0.19; 50.0 ms: 0.51 ± 0.29), while white boxplots represent the corresponding null distributions (12.5 ms: 0.33 ± 0.12; 5.0 ms: 0.61 ± 0.15; 50.0 ms: 0.38 ± 0.26). For all IPIs, the model significantly outperformed the shuffled baseline (Wilcoxon signed-rank test; 12.5 ms: p < 0.001; 25.0 ms: p = 0.010; 50.0 ms: p < 0.001). **d)** Control accuracy as a function of IPI and the number of stimulated channels (*κ*) at each timestep. A linear mixed-effects model with random intercepts for each session revealed a significant quadratic relationship between the number of stimulated channels and control accuracy (*κ*: *β* = 1.077, p = 0.014; *κ*^2^: *β* = –0.264, p = 0.006). The IPI itself (*β* = –0.012, p = 0.278) and its interactions with channel count (all p > 0.18) did not significantly modulate performance. The black line indicates the population-level quadratic fit. **e)** Spatial independence of control accuracy. Control accuracy as a function of the average Euclidean distance between stimulating elecrodes. This analysis was restricted to stimulation sequences involving *κ* = 2 channels per timestep, as in the main results, to isolate spatial effects. A linear mixed-effect model indicated that electrode distance was not a significant predictor of accuracy (*β* = 0.001, p = 0.933), suggesting that the control framework is robust to the spatial arrangement of stimulation sites.

**FIG. S3.**
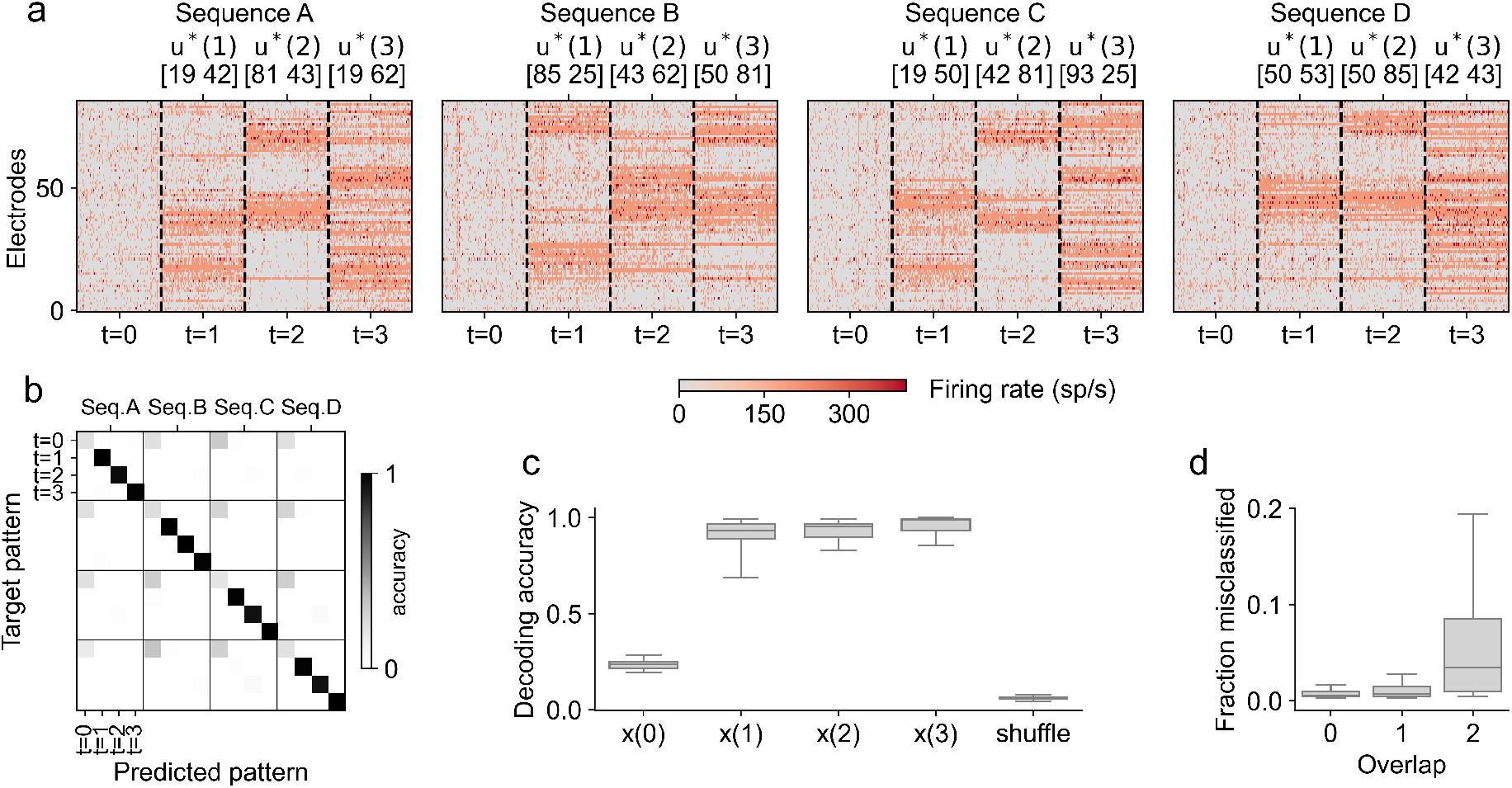
Neural population state discrimination across stimulation sequences. **a)** Single-trial evoked activity for four different sequences. Each sequence evokes distinct spatiotemporal population dynamics that are consistent across trials. The identity of stimulated electrodes at each time step is shown on top. Each sequence was repeated 100 times; repetitions are concatenated horizontally within each time step. **b)** Sequence and time-step discrimination visualized through a confusion matrix for a linear discriminant analysis (LDA) classifier. High values on the diagonal represent accurate simultaneous discrimination of both the stimulation sequence and the time step within that sequence. At t = 0, the absence of a clear diagonal structure indicates that initial conditions are not decodable between sequences, confirming that discrimination is driven by the stimulation patterns rather than pre-existing state differences. **c)** Summary of decoding accuracy across all sessions compared to randomized shuffle distributions for each timestep t in the sequence (x(0): 0.24 ± 0.03, mean ± s.d.; x(1): 0.91 ± 0.08; x(2): 0.93 ± 0.05; x(3): 0.96 ± 0.04; shuffle: 0.06 ± 0.01; naive chance level, 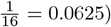. Although decoding accuracy for x(T ≥ 1) was close to perfect (1.0), a linear mixed-effect model revealed a modest but significant increase in decoding accuracy over time (*β* = 0.028, p < 0.001), consistent with a cumulative effect of sequential stimulations on the discriminability of the evoked population activity. **d)** The impact of stimulation overlap on misclassification, shown as the fraction of misclassified patterns as a function of the number of overlapping stimulation sites between sequences. Low error rates across all conditions demonstrate that population dynamics remain discriminable despite shared stimulation electrodes (Kruskal-Wallis: H = 155, p < 10^−10^).

**FIG. S4.**
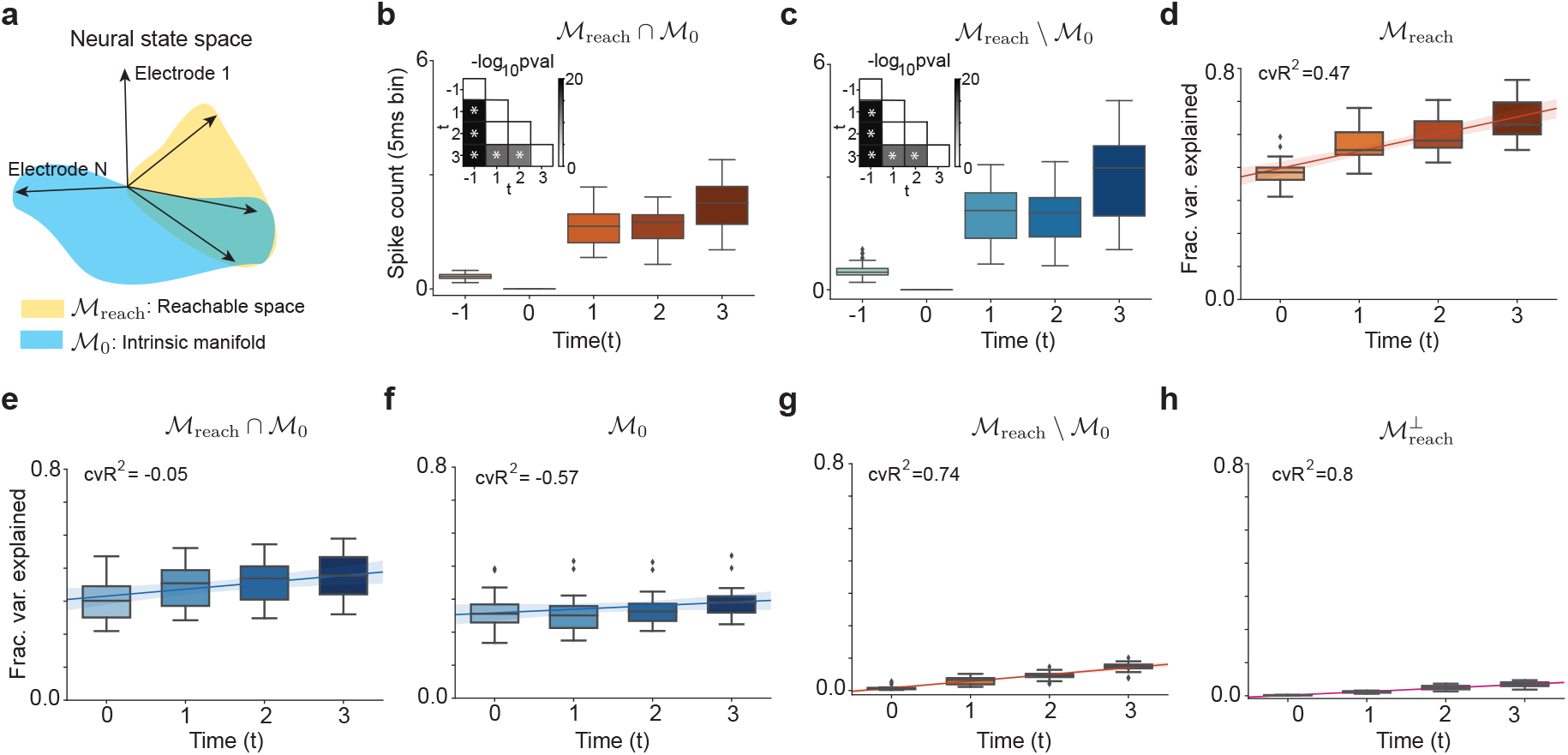
Traversability within shared and non-shared components of the reachable and intrinsic manifolds. **a)** Schematic of the neural state space, using the same notations as in Fig. 4. The reachable manifold ℳ_reach_, the intrinsic manifold ℳ_0_, and their overlap ℳ_reach_∩ℳ_0_ are illustrated. **b)** Distance between the pre-stimulation state x(0) and the evoked activity x(t) when trajectories are projected onto the subspace of the reachable manifold shared with the intrinsic manifold, ℳ_reach_ ∩ ℳ_0_. **c)** Distance between x(0) and x(t) when trajectories are projected onto the subspace of the reachable manifold that is not shared with the intrinsic manifold, ℳ_reach_\ℳ_0_. **d-h)** Fraction of variance explained for different manifolds along the neural trajectories X(0:t) as a function of time t for all stimulation patterns in each session, where X(0:t) denotes the collection of population activity states from t = 0 up to t. **d)** Fraction of variance explained of the reachable manifold ℳ_reach_.**e)** Fraction of variance explained of the reachable manifold shared with the intrinsic manifold. **f)** Fraction of variance explained of intrinsic manifold X_0_. **g)** Fraction of variance explained of the reachable manifold not shared with the intrinsic manifold. **h)** Fraction of variance explained in the full activity space that lies outside the reachable manifold, 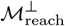. Lines show a linear fit. Shading indicates 95% confidence interval.

**FIG. S5.**
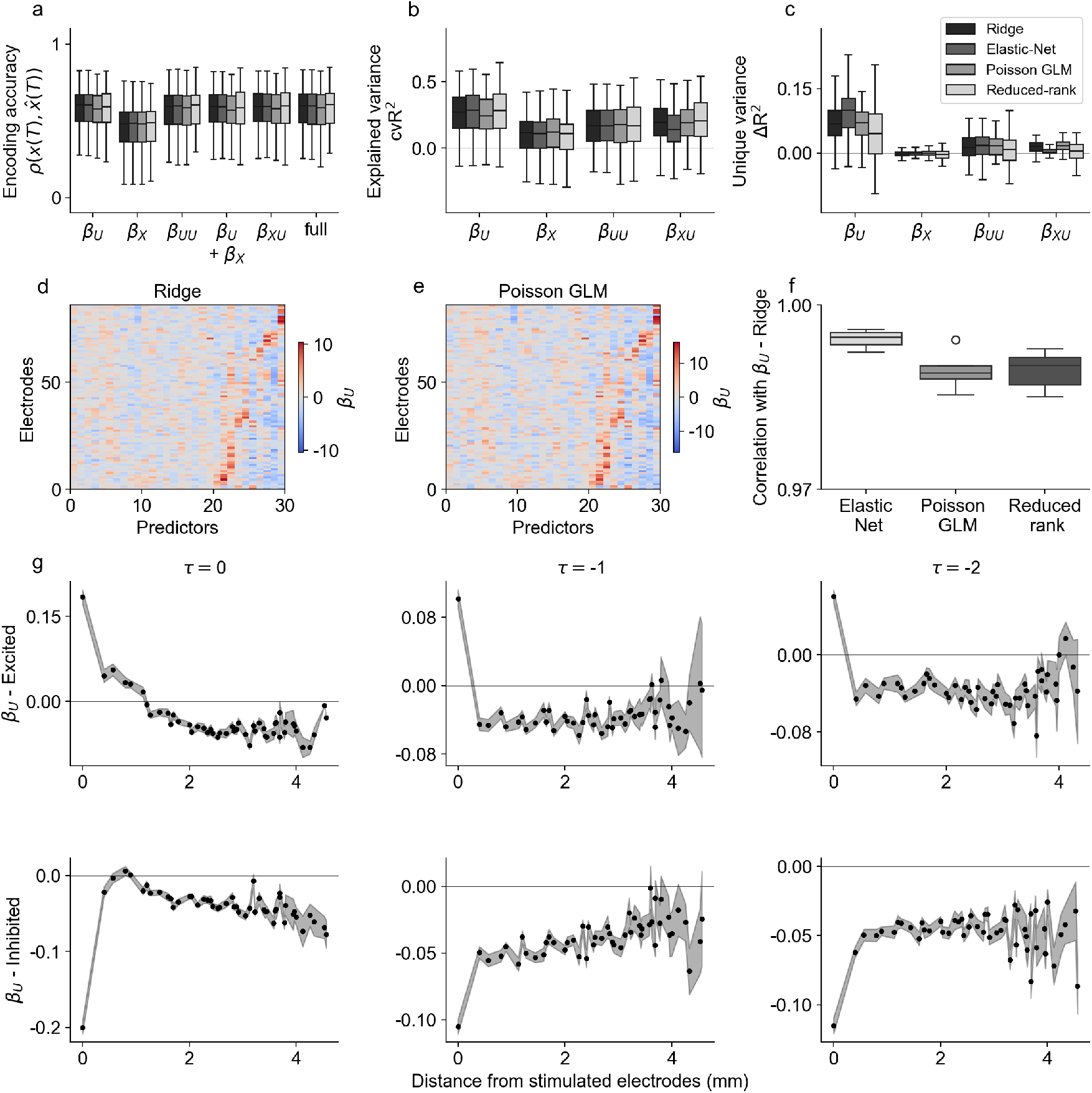
Performance and validation of the encoding models. **a)** Encoding accuracy of different model components. Comparison of prediction accuracy (*ρ*) for the full encoding model (Eq. 20) and reduced models with specific predictor sets: stimulation history (*β*_U_), spike history (*β*_X_), and interaction terms (*β*_UU_, *β*_XU_). **b)** Cross-validated explained variance (cvR^2^) for individual predictor sets across sessions, quantifying the total variance each set can explain when considered alone. **c)** Unique variance (ΔR^2^) contributed by each predictor set. Unique variance is computed by comparing the full model against shuffle-in models where one predictor set is shuffled across trials while all other predictors are left intact. The stimulus-driven component (*β*_U_) explains the largest fraction of unique variance across sessions. **d-e)** Comparison of parameter estimation methods. Visualization of estimated stimulus-response coefficients (*β*_U_) across all measured electrodes using Ridge regression (d) and Poisson GLM (e). The close correspondence between the estimated stimulation fields (*β*_U_) indicates that the linear Gaussian approximation effectively captures the dominant stimulation-driven structure. In the example session shown, the number of predictors in the *β*_U_ set equals the number of stimulated electrodes (m = 10) times the number of pulses in the stimulation sequence (T = 3); predictors 20-30 correspond to the last pulse. **f)** Robustness across regularization and noise models. Correlation coefficients between the Ridge-derived *β*_U_ parameters and those derived from Elastic-net, Poisson GLM, and reduced-rank models. High correlation values (> 0.98) indicate that the identified stimulation fields are robust to the choice of regularization and observation model. **g)** Spatial decay and temporal evolution of stimulation-driven influence. Interaction kernels plotted as a function of Euclidean distance from the stimulated electrodes at different time lags (τ = 0, –1, –2). Primary excitatory (top) and inhibitory (bottom) influences show a characteristic decay with distance, and a temporally decaying spatial footprint of microstimulation effects. Gray shading indicates s.e. across electrodes.

